# Intelligent Resolution: Integrating Cryo-EM with AI-driven Multi-resolution Simulations to Observe the SARS-CoV-2 Replication-Transcription Machinery in Action

**DOI:** 10.1101/2021.10.09.463779

**Authors:** Anda Trifan, Defne Gorgun, Zongyi Li, Alexander Brace, Maxim Zvyagin, Heng Ma, Austin Clyde, David Clark, Michael Salim, David J. Hardy, Tom Burnley, Lei Huang, John McCalpin, Murali Emani, Hyenseung Yoo, Junqi Yin, Aristeidis Tsaris, Vishal Subbiah, Tanveer Raza, Jessica Liu, Noah Trebesch, Geoffrey Wells, Venkatesh Mysore, Thomas Gibbs, James Phillips, S. Chakra Chennubhotla, Ian Foster, Rick Stevens, Anima Anandkumar, Venkatram Vishwanath, John E. Stone, Emad Tajkhorshid, Sarah A. Harris, Arvind Ramanathan

## Abstract

The severe acute respiratory syndrome coronavirus-2 (SARS-CoV-2) replication transcription complex (RTC) is a multi-domain protein responsible for replicating and transcribing the viral mRNA inside a human cell. Attacking RTC function with pharmaceutical compounds is a pathway to treating COVID-19. Conventional tools, e.g., cryo-electron microscopy and all-atom molecular dynamics (AAMD), do not provide sufficiently high resolution or timescale to capture important dynamics of this molecular machine. Consequently, we develop an innovative workflow that bridges the gap between these resolutions, using mesoscale fluctuating finite element analysis (FFEA) continuum simulations and a hierarchy of AI-methods that continually learn and infer features for maintaining consistency between AAMD and FFEA simulations. We leverage a multi-site distributed workflow manager to orchestrate AI, FFEA, and AAMD jobs, providing optimal resource utilization across HPC centers. Our study provides unprecedented access to study the SARS-CoV-2 RTC machinery, while providing general capability for AI-enabled multi-resolution simulations at scale.

## 1 JUSTIFICATION

We developed an AI-enabled multi-resolution simulation framework for studying complex biomolecular machines by directly integrating experimental data. Our framework sets high-water marks for AI-driven multi-resolution simulations and achieving high utilization of resources across diverse supercomputing platforms at multiple sites.

## 2 PERFORMANCEATTRIBUTES

2 PERFORMANCE ATTRIBUTES
| Performance Attribute | Our Submission |
| --- | --- |
| Category of achievement | Scalability, Time-to-solution |
| Type of method used | Explicit, Deep Learning |
| Results reported on the basis of | Whole application including I/O |
| Precision reported | Mixed Precision |
| System scale | Measured on full system |
| Measurement mechanism | Hardware performance counters, Application timers, Performance Modeling |

## 3 OVERVIEW OF THE PROBLEM

The novel coronavirus 2019 (COVID-19) pandemic has led to a massive acceleration in the pace of development in experimental structural biology. Multiple research teams have invested their best resources towards the common goal of characterizing viral components and their biological functions, thereby providing a sign-post for the future direction of the field (Alam and Higgins, 2020, Bárcena et al., 2021, Barrantes, 2021, Kim and Jung, 2021). Structural biology data, which provides the basis for rational design of all new medicines against human and animal disease, is now inherently multi-modal and multi-scale, requiring novel integrative approaches (AlQuraishi, 2019, Arantes et al., 2020, Jumper et al., 2021, Minkyung et al., 2021, Muratov et al., 2021, Padhi et al., 2021, Tunyasuvunakool et al., 2021, Zimmerman et al., 2020).

While the mechanism of human host cell entry and infection by SARS-CoV-2 via the spike glycoprotein is now relatively well characterized (Barros et al., 2020, Shang et al., 2020, Sztain et al., 2021, Zhang et al., 2021), how the SARS-CoV-2 replicates inside the host cell is still unclear (Romano et al., 2020). The viral-RNA replication mechanism is complex, involving RNA synthesis, proofreading, and capping and is mainly carried out by the mini-replication transcription complex plus error correction machinery (mRTC+ECM from here on referred to as RTC), having to survive against the human immune response. Cryo-EM techniques and computational methods have been immensely helpful in elucidating the overall structural organization of the RTC (Chen et al., 2020, Perry et al., 2021, Yan et al., 2020, 2021a), but the high intrinsic flexibility, size and complexity of the nsp arrangement entails that the overall resolution of the data is inherently poor. Consequently, the structure refinement workflows discard 30-40% of collected images from the existing RTC complex datasets (Chen et al., 2020, Yan et al., 2020, 2021a), leaving significant gaps in our understanding.

Although several studies have focused on disrupting the function of the individual non-structural proteins (nsps) with small molecules, key insights into the overall structural organization, dynamics and function of the RTC machinery are more difficult to obtain. This is crucial, because the ability to target protein-protein interactions between subunits of the RTC complex offers far more possibilities for drug development. Moreover, the arrangement of individual protein components of the RTC+ECM protein is itself dynamic during the viral life-cycle. The computational capability to model these interactions would provide further insight of relevance to drug development, but is currently impossible without novel multi-scale models and the workflows that connect them.

The primary challenge of experimental imaging is elucidating diverse structural dynamics. This stems from the *averaging* process of the imaging data: cryo-EM, in particular, and other experimental techniques capture only the most sampled conformational states as static, snapshot-like representations, but the intermediates or transitional states are less represented (Lyumkis, 2019, Merk et al., 2016). The details of motions within flexible domains can be enriched using complimentary tools such as molecular dynamics (MD) simulations and Bayesian inference techniques (Bowerman et al., 2017, Bratholm et al., 2015, Cavalli et al., 2007, Grishaev and Llinás, 2005, Scheres, 2012); however, the timescales accessible to these atomistic simulations can be a limiting factor. In addition, advances in 4D imaging modalities (Earnest et al., 2017, Engel et al., 2015, Mahamid et al., 2016, Villa and Lasker, 2014) and the volume of data generated from such experimental datasets can be overwhelming.

Therefore, in this paper we address the urgent, yet unmet need to develop scalable computational tools that can aid the improvement of resolution within cryo-EM datasets through multi-resolution simulations. In an effort to bridge the gap between experimental and purely all-atom molecular dynamics (AAMD), we leverage a complementary mesoscale method of representing biophysical systems, treating biomolecules as visco-elastic continuum solids using fluctuating finite element analysis (FFEA) (Oliver et al., 2013) technique. These continuum-scale lower resolution FFEA simulations provide a generative model for the cryo-EM data. However, implementing an approach that directly models electron density information from cryo-EM data requires a radically different way to model conformational ensembles, one that moves away from atomistic-resolution towards a *continuum*-representation, where by the intrinsic resolution of the data can be captured with nodes and meshes, allowing experimentally obtained cryo-EM data to be used directly in simulations, at time and length-scale orders of magnitude increased over traditional all-atom methodologies.

We present a radically innovative workflow, providing the opportunity to accelerate conformational sampling (both AAMD and FFEA) with AI, thus holding much promise in providing mechanistic insights into the dynamics of large biomolecular complexes (Fig. 1). We leverage FFEA and AAMD simulations to iteratively fill in the gaps of cryo-EM data by refining the pairwise atomic/molecular interactions while being automatically constrained by the degrees of freedom embodied in the cryo-EM data. This iterative coarse-graining approach maintains the fidelity between experimentally observed densities with the detailed potentials from rigorous physics-based AAMD simulations to learn informative *priors* and effectively guide the conformational search and can result in better fits to the experimental observations. We demonstrate our iterative approach in modeling the complex conformational transitions within the SARS-CoV-2 RTC to observe how the various nsps coordinate the proofreading process of the viral-RNA. These conformational changes can aid in understanding and guiding the design of novel therapeutics such as molnupiravir, which aim to target specific nsps. (Agostini et al., 2019, Sheahan et al., 2020)

**Figure 1:**
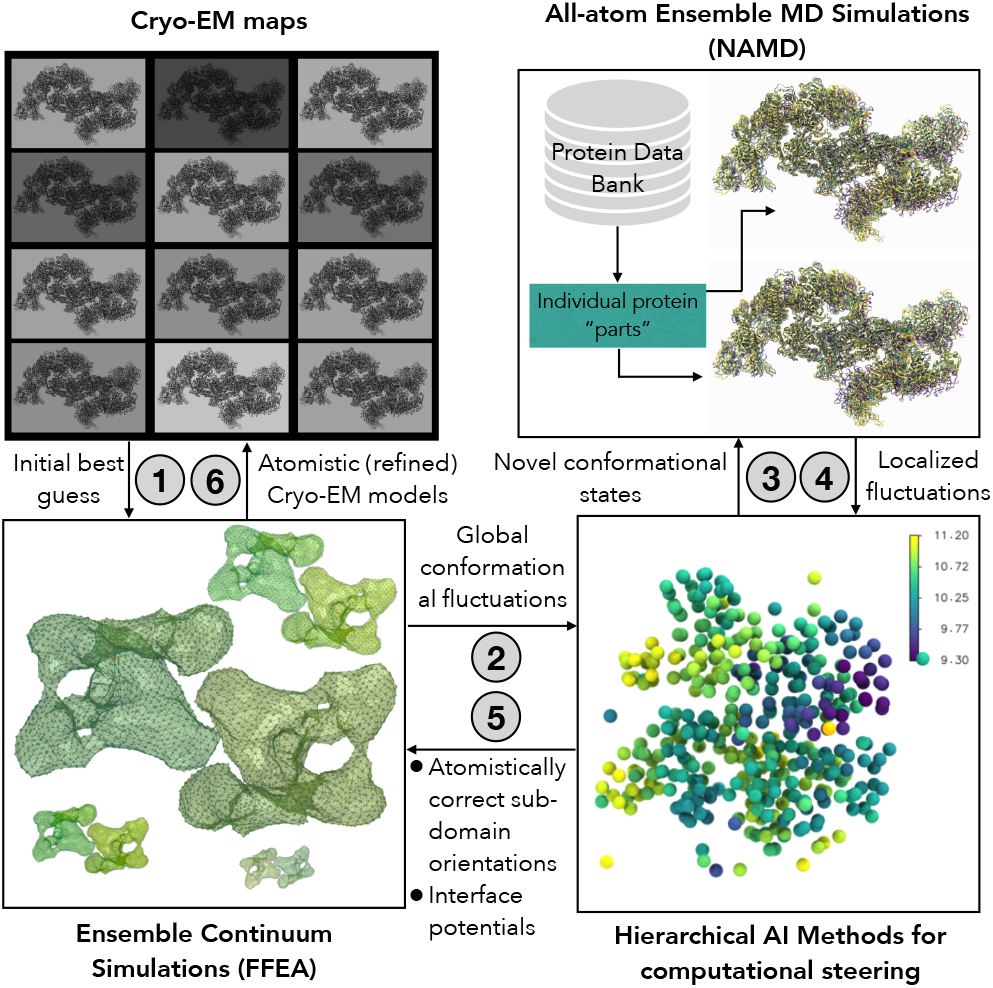
An integrative biology framework for refining low resolution cryo-EM structures with multi-resolution simulations. (1) Representing the cryo-EM density map as a continuum visco-elastic solid. (2) Finite element analysis simulations are then used to generate new conformations. AI techniques identify interesting events in the landscape (global conformational changes), while (3) simultaneously constraining them with all-atom simulations derived protein-protein interface potentials. (4) AI methods are also used to learn local conformational changes across the molecular machine, such that they can be used to (5) refine domain orientations in the entire biomolecular complex. (6) The output represents a set of atomistically refined ensemble of structures that captures the conformational fluctuations embodied in the cryo-EM data.

**Figure 2:**
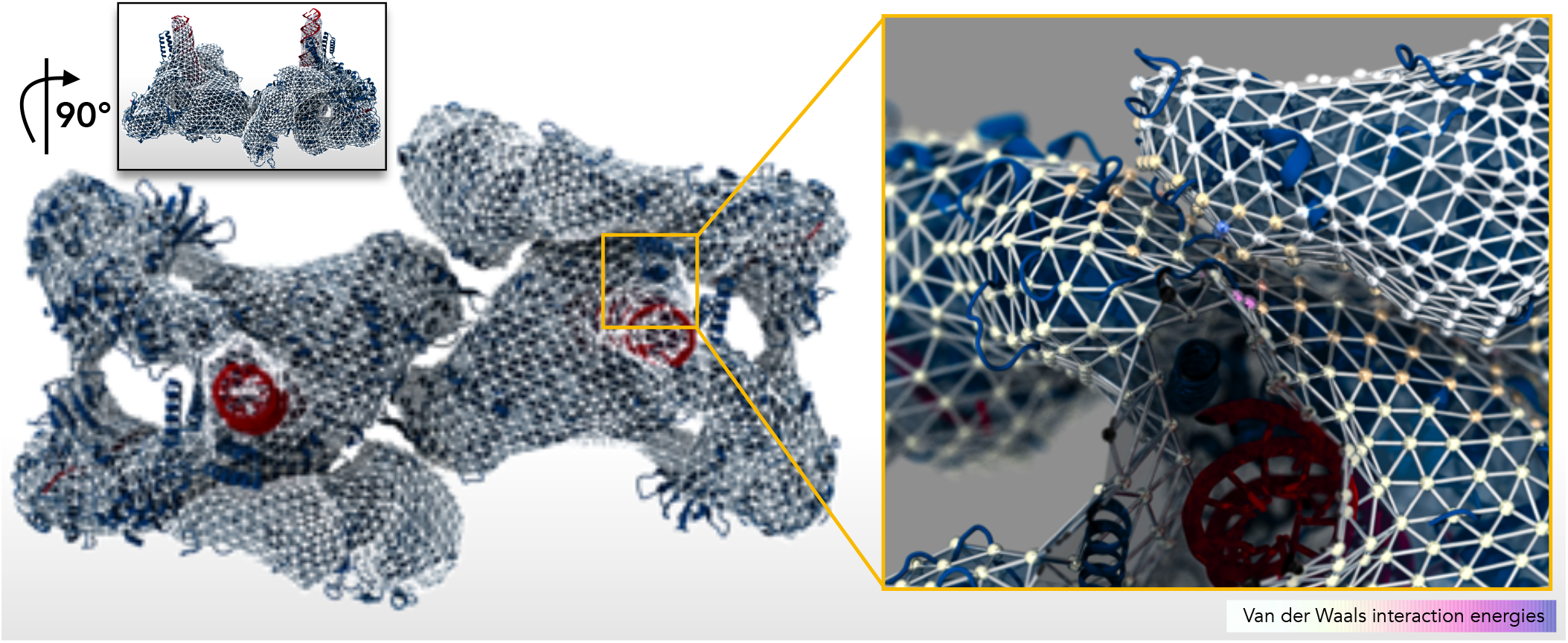
Hybrid structure of the FFEA mesh superimposed with the all-atom representation. The all-atom structure of the RTC dimer is shown as a cartoon (blue) and the FFEA tetrahedral mesh structure determined from the experimental cryo-EM map is shown as a wireframe. The top inset represents a 90° rotation of the RTC dimer capturing the range of protein-protein interfaces in the machine. A close-up view of the mesh at the interface between the RdRp (nsp12) and nsp13 reveals the surface interface potentials inferred from AAMD simulations. Each mesh point is painted with the surface interface potentials, with darker shades of red indicating higher and lighter shades indicating lower interaction energy in the protein-protein interface.

## 4 CURRENT STATE OF THE ART

### 4.1 Parallel molecular dynamics

NAMD (Phillips et al., 2005, 2020) has been one of the most utilized parallel molecular dynamics engines for over two decades, being cited once every ~70 minutes ^1^. NAMD uses adaptive, asynchronous, message-driven execution based on Charm++ (Kalé et al., 2019, Kalé and Zheng, 2013), while efficiently utilizing GPUs (Phillips et al., 2008). It contains advanced features giving scientists an extensive set of tools to observe and/or bias their systems using state-of-the-art simulation methodologies such as collective variables (Fiorin et al., 2013) and molecular dynamics flexible fitting (MDFF) (Trabuco et al., 2009, Vant et al., 2020) modules.

VMD and its psfgen plugin were used to build and analyze the molecular complexes studied herein. Simulation preparation, visualization, and conventional analysis approaches were performed using VMD, with extensive use of GPU-accelerated remote visualization resources (Humphrey et al., 1996a, Stone et al., 2013a,b, 2016). VMD incorporates features for processing cryo-EM density maps to prepare and analyze *mdff* hybrid fitting simulations as required in this project (Stone et al., 2014). VMD incorporates a custom GPU-accelerated ray tracing engine that exploits hardware accelerated construction of ray tracing BVH acceleration structures, BVH traversal, and ray-triangle intersections (Sener et al., 2021).

### 4.2 FFEA: continuum molecular simulations

Recent advances in cryo-EM, and -ET aided scientists to generate structural and dynamic information of large biomolecules in 3D volumetric data (Kühlbrandt, 2014). Expanding on this volumetric shape, FFEA uses a 3D tetrahedral finite element mesh providing dynamic insight of the structure alone and in response to interactions with other molecules (Solernou et al., 2018). FFEA is a new physics algorithm for simulation of mesoscale biological structures obtained from cryo-EM and cryo-ET (Solernou et al., 2018). FFEA treats biomolecules as continuum visco-elastic solids subject to thermal noise, chosen so as to satisfy the fluctuation-dissipation theorem (Oliver et al., 2013). To describe interacting proteins, FFEA includes short-range van der Waals attractive forces and steric repulsion. Therefore, FFEA uses a unique “top-down” rather than the more conventional “bottom-up” coarse-graining strategy, by introducing nanoscale thermal fluctuations into macroscopic continuum mechanics equations.

FFEA also includes functionality to exert external forces on proteins, to connect proteins together with harmonic springs and to represent conformational changes between distinct protein conformational states, for example between the pre- and post-powerstroke states of molecular motors (Richardson et al., 2020). FFEA has been used to successfully model diverse biological systems, including the rotary ATPase motor (Richardson et al., 2014), axonemal (Richardson et al., 2020) and cytoplasmic (Hanson et al., 2021) dynein motors, and protein antibodies subjected to external forces (van der Heijden et al., 2020).

### 4.3 Multiscale biological simulations

Integrating data across diverse spatial, temporal, and functional scales has played an important role in understanding the role of molecular interactions in various diseases (Pak and Voth, 2018, Tozzini, 2010, Walpole et al., 2013, Zhou, 2014). In order to bridge information across multiple scales, a number of coarse-graining approaches have been widely employed. These methods have been used to study individual proteins from the SARS-CoV-2 proteome (De Sancho et al., 2020, Garay et al., 2021), as well as more complex systems such as the entire SARS-CoV-2 virion (Yu et al., 2021). We note that compared to the state-of-the-art approaches, we choose a continuum representation of molecules, one that is more conducible for accessing much larger length- and time-scales using FFEA. While finite element methods are used to model biological systems (e.g., growth models, bone growth, etc.) (Dong and Skalak, 1992, Kennaway and Coen, 2019, Panagiotopoulou et al., 2017), we note that our collective approach enables tight integration between all-atom and continuum representations for large biomolecular complexes such as the SARS-CoV-2 RTC.

While physics-based approaches for coarse-graining are usually employed to overcome limitations of the space- and time-scales accessed by traditional molecular simulations, machine learning techniques are now being regularly employed to adaptive coarsegrained simulations from all-atom simulations (Durumeric and Voth, 2019, Husic et al., 2020, Noé, 2020).

### 4.4 AI enabled adaptive MD simulations

MD simulation provides atomistic details of molecular interactions and ensued dynamics. It’s however computationally expensive, and the sampling efficiency can be severely hindered, when trapped by local minima in the free energy landscape. Our group has developed AI-enabled adaptive MD simulations, namely DeepDriveMD (Lee et al., 2019), which records MD propagation in latent space via convolutional variational autoencoder (CVAE) (Bhowmik et al., 2018), and drives adaptive sampling in the high dimensional conformational landscape. The AI inference constantly observes the simulation runs, prunes stagnant ones that are trapped in local energy minima, and spawns new ones from less sampled conformations. We have shown that this approach can accelerate sampling of rare events (for example, in the SARS-CoV-2 Spike opening simulations (Casalino et al., 2021), protein folding (Lee et al., 2019)) by at least an order of magnitude. When integrated with specialized AI-hardware to accelerate the learning, it can provide nearly 4 orders of magnitude speedup (Brace et al., 2021).

### 4.5 Workflow infrastructure

Distributed science workflows must invariably handle data and control flow across networks spanning administrative domains. Challenging barriers to workflow management frameworks’ adoption in multi-user HPC environments lie in deploying distributed client/server infrastructures and establishing connectivity among remote systems. For instance, Fireworks (Jain et al., 2015) and the RADICAL-Pilot / Ensemble Toolkit (EnTK) (Balasubramanian et al., 2016, Merzky et al., 2018) are three widely-used WMFs at supercomputing facilities that expose Python APIs to define and submit directed acyclic graphs (DAGs) of stateful tasks to a database. These WMFs possess various implementations of a common *pilot job* design, whereby tasks are efficiently executed on HPC resources by a process that synchronizes state with the workflow database. Because the database is directly written by user or pilot job clients, users of these WMF typically deploy and manage their own database servers such as MongoDB servers.^2^ These methods are non-portable and depend on factors such as whether the compute facility mandates multi-factor authentication (MFA).

In the context of REST interfaces to HPC resources, efforts such as the *Superfacility* concept (Enders et al., 2020) envisions a future of automated experimental and observational science workflows linked to national computing resources through Web APIs. Implementations such as the NERSC Superfacility API^3^ and the Swiss National Supercomputing Centre’s FirecREST API (Cruz and Martinasso, 2019) expose methods to submit and monitor batch jobs, move data between remote filesystems, and check system availability. However, these facility services alone do not address workflow management or high-throughput execution; instead, they provide web-interfaced abstractions of the underlying facility, analogous to modern cloud storage and infrastructure services.

funcX (Chard et al., 2020) is an HPC-oriented instantiation of the function-as-a-service (FaaS) paradigm, where users invoke Python functions in remote containers via an API web service. funcX endpoints run on the login nodes of target HPC resources where Globus Auth is used for authentication and user-endpoint association. The funcX model is tightly focused on Python functions and therefore advocates decomposing workflows into functional building blocks. Each function executes in a containerized worker process running on one of the HPC compute nodes. This fails to support a generalized model of applications such as simulations with executables that may or may not leverage containers, per-task remote data dependencies, programmable error- and timeout-handling, and flexible per-task resource requirements (e.g. tasks may occupy a single core or specify a multi-node, distributed memory job with some number of GPU accelerators per MPI rank).

Balsam (Salim et al., 2021) provides a unified API for describing distributed workflows, and the user site agents manage the full lifecycle of data and control flow. The Balsam service-oriented architecture shifts administrative burdens away from individual researchers by routing all user, user agent, and pilot job client interactions through a hosted, multi-tenant web service. As Balsam execution sites communicate with the central service only as HTTP clients, deployment involves a simple user-space pip package installation on any platform with a modern Python and outbound internet access. Due to the ubiquity of HTTP data transport, Balsam works “out of the box” on supercomputers spanning DOE and NSF facilities against a cloud-hosted Balsam service.

## 5 INNOVATIONS REALIZED

Our innovations can be briefly summarized as follows. (1) We implement an automated workflow to seamlessly transition from AAMD to FFEA via an AI-based approach. By optimizing the performance of AAMD simulations (Sec. 5.1) we achieve optimal performance on modern multi-GPU-based computing systems. (2) By clustering the conformations from AAMD simulations using unsupervised AI methods (Sec. 5.3), we learn the parameters for constraining the FFEA simulations based on the implicit fluctuations between protein-protein interfaces inferred from AAMD simulations. We develop a novel AI-approach, namely Graph neural operators (GNO) (Sec. 5.3.3) to summarize time-dependent conformational changes observed from ensemble AAMD simulations. (3) To overcome the bottleneck of training deep learning models on large volumes of data generated by AAMD simulations, we scaled their performance on emerging AI-hardware such as the Cerebras CS-2 accelerator as well as on the Summit supercomputer.^4^ (4), Finally, to our knowledge, this is the first attempt at executing a single coordinated workflow across multiple supercomputing facilities. We achieved this through the Balsam multisite workflow manager (Sec. 5.4). The performance gains enabled by these innovations provide a generalized, multi-scale computational toolkit for exploring dynamic biomolecular machines.

### 5.1 NAMD all-atom MD simulations

#### 5.1.1 GPU-resident NAMD 3 for GPU-accelerated MD simulations

To achieve previously unrealized performance levels for MD simulations on GPU-accelerated HPC platforms, we have developed a major new “GPU-resident” computing approach, implemented in NAMD 3. The GPU-resident computing mode in NAMD 3 employs GPU acceleration for the most common molecular dynamics force and energy calculations together with time integration and rigid bond constraints, allowing sequences of timesteps to be simulated entirely on the GPU without any significant host–GPU data transfers, thereby exploiting a significantly higher fraction of theoretical peak GPU performance (Phillips et al., 2020). Most recently, we have further extended the GPU-resident capabilities of NAMD 3 to enable strong-scaling of a simulation across multiple GPUs, with particular emphasis on hardware platforms that incorporate NVLink connectivity among peer GPUs. NVLink facilitates direct load/store memory access among peered GPUs with low latency and high bandwidth data exchange performance levels far beyond what is currently possible with a conventional HPC network interconnect, or even host-internal PCIe transfers among GPUs.

The multi-GPU simulation algorithm generally mirrors the techniques used in the conventional distributed-memory builds of NAMD, except that it is designed to leverage the completely unified virtual memory address space of the GPUs to exchange data directly through fine-grained memory load/store operations. NVLink support for direct fine-grained memory loads/stores in concert with atomic increment and atomic compare-swap operations permit efficient implementation of performance-critical multi-GPU reductions and synchronization of GPUs within algorithm phases. Platforms providing GPUs with a fully connected topology permit strong scaling of molecular systems ~1M atoms on up to 8 GPUs with acceptable efficiency, limited primarily by poor scalability of the PME full electrostatics algorithm.

While running a simulation per GPU avoids these scaling challenges, multi-GPU scaling enables simulation over longer timescales. For systems of ~1M atoms, the timescale is typically a limiting factor to study conformational changes that a protein undergoes, essential in understanding how a complex functions. Performance considerations dictate that any advanced sampling method implemented for GPU-resident must avoid frequent data transfers to and from the host. Many of these features are contributed by external methodological contributors and have not yet been implemented to run on GPUs. As such, NAMD 2 and CPU-based resources still play an important role in the workflow.

#### 5.1.2 VMD full-time interactive ray tracing

To facilitate creation of high-fidelity visualizations with high interactivity, VMD has been extended with a new real-time ray tracing (RTRT) rendering mode that uses full-time interactive progressive refinement ray tracing for rendering of the molecular scenes instead of OpenGL rasterization and includes ambient occlusion lighting, shadows, high-quality transparency, and depth of field focal blur. The new RTRT rendering mode provides substantially more visually informative visualizations of complex molecular scenes as compared with conventional OpenGL rasterization, by bringing offline-quality rendering features to the interactive VMD display window for the first time. When the RTRT rendering mode is active, advanced visualization features that were formerly only usable for offline rendering, or for static structures, can now be applied to simulation trajectories while maintaining full interactivity.

#### 5.1.3 RTC System Preparation for all-atom simulations with NAMD

The SARS-CoV-2 RTC is a multi-subunit structure composed of several nsps, including the RNA-dependent RNA polymerase (RdRp, nsp12), Zinc-bound helicase (HEL, nsp13), the RNA-capping enzymes such as the nsp14 (N7-methyltransferase/MTase) and nsp16 (O2-MTase), the proofreading enzyme, namely nsp14, and uridylate-specific endoribonuclease activity (NendoU, nsp15)(Fig. 3A) (Chen et al., 2020, Perry et al., 2021, Yan et al., 2020, 2021a). Together they participate in the process of reproducing the viral genome (Wu et al., 2020). This complex plays an important role in the viral life cycle, and is therefore an important drug target (Gao et al., 2020, Wang et al., 2020). Revealing the atomistic details of this complex is essential for understanding how the viral RNA is processed.

**Figure 3:**
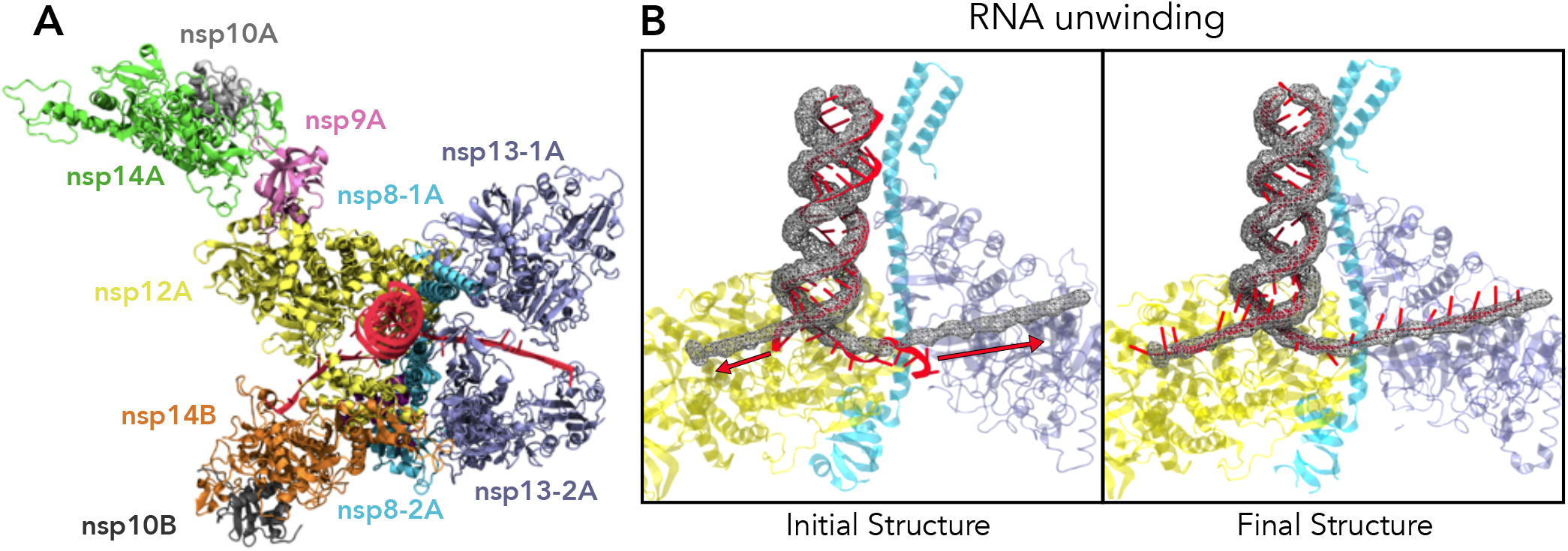
The atomistic representation of a functional RTC monomer (A) with the RNA (red), extended to the active site of the other nsp14-10 unit of the opposite monomer (nsp14B - orange), and the exit site on the helicase of the same monomer (nsp13-2A). The mechanism of RNA unwinding using non-equilibrium molecular dynamics simulations starting from the initial structure as captured in 7EGQ (B-left), to the final structure (B-right). The RNA backbone is restrained with an *mdff grid*, represented in a grey mesh, while using *colvars* to pull the strands to the active sites on the protein. This is an unprecedented simulation of the mechanism of RNA unwinding which can give significant insight into the RTC function.

**Figure 4:**
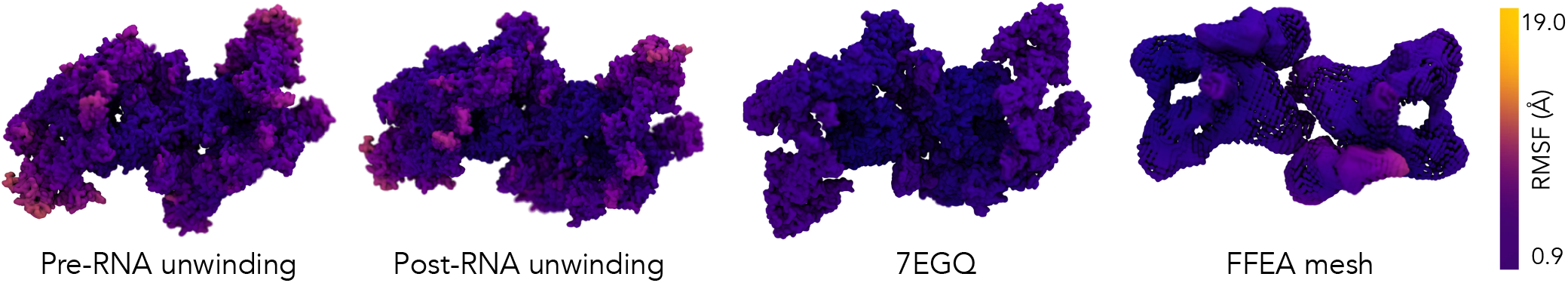
Root-mean squared fluctuations (RMSF) of the SARS-CoV-2 RTC provide insights are helpful in determining the intrinsic concerted motions in the RTC that are implicitly encoded in the experimental data. The structure of the RTC is painted using the RMSF determined from the ensemble AAMD simulations labeled pre-RNA-unwinding and post-RNA-unwinding states, the cryo-EM structure (PDB id: 7EGQ; where we use the experimental B-factors to approximate the RMSF) and FFEA simulations. While the FFEA simulations show larger fluctuations at the interface of nsp10-14 as it interacts with the two nsp13 subunits, both the AAMD simulations capture larger fluctuations in the two nsp13 subunits across each monomer. While this embodies slightly different rearrangements between the FFEA and AAMD simulations, the relative fluctuations across these subunits are high within the cryo-EM data indicating the dynamic nature of this assembly.

We prepared the structure of the Cryo-EM dimeric form of the complex 7EGQ.pdb (Yan et al., 2021b). Cations have functional significance for the RTC, therefore the linking HIS and CYS residues were mutated to HSE/HSD and CYM respectively, to have the correct ionization state to coordinate the present 3 Mg^2+^ ions and 26 Zn^2+^ ions. A complementary CYT base was added to the end of two incomplete RNA strands. Missing loop residues ARG ALA ARG corresponding to residue numbers 314-316 in the nsp13 subunits were modeled using pdbfixer tool in OpenMM (Eastman et al., 2017). The protein subunits were capped with an NTER and CTER patches. The psfgen package from VMD (Visual Molecular Dynamics) (Humphrey et al., 1996b) was used to assemble the system, which was then solvated using the SOLVATE plugin and neutralized with K^+^ and Cl^-^ ions for a concentration of 150 mM KCl using AUTOIONIZE. The resulting systems consist of ~1.1 M atoms and box size of ~ 168 × 220 × 312Å^3^. The systems were energy minimized using steepest descent method for 500 steps, followed by a 3 ns equilibration protocol restraining all but the water molecules using a constant k = 3 kcal mol^−1^ Å^−2^. The system was further equilibrated restraining only the protein backbone alpha carbon atoms and nucleic backbone phosphate atoms with a constant of k = 0.05 kcal mol^−1^ Å^−2^ with NAMD2.14 (Phillips et al., 2005) NVT ensemble using CHARMM36 forcefields for proteins and nucleic acids (Vanommeslaeghe et al., 2010).

#### 5.1.4 Grid-steered Molecular Dynamics Simulation of RNA Unwinding

One of the crucial functions of the RTC complex is to proofread the RNA, which is a process that requires a major conformational change of the protein complex and unzipping of the RNA. To mimic this operation, we performed extensive preliminary studies using an isolated RNA double-helical structure from the aforementioned prepared structure in order to optimize the combination of grid-steered molecular dynamics (GMD) (Wells et al., 2007) and collective variables (colvars) (Fiorin et al., 2013) and their respective optimal force constants to mimic the corkscrew motion of the RNA unwinding. To speed up these empirical studies, instead of the all-atom representation, the protein was represented by a molecular dynamics flexible fitting *mdff* grid map exerting a repulsive *gridForce* (Trabuco et al., 2009). A variety of *colvars* were employed to drive the extension of the RNA structure, including *distanceZ* to pull on the strands to the nsp14-10 active site and the exit point on the nsp13 helicase, as proposed in previous studies (Yan et al., 2021b). We applied a rotational force by using the *orientationAngle* colvar, while also pulling on the center of mass using *distance* colvar. The final conformation used for the simulations are a hybrid of the initial double-helical structure of the RNA strands, with the extended single strands which have reached their target points within the complex (Fig. 3B). This structure was obtained by building an attractive 3 dimensional electron density map manually constructed from the backbone of the initial RNA position with the final extended strands obtained from the non-equilibrium simulations.

The all-atom simulations to mimic this process were started from the equilibrated structure described in section 5.1.3. We performed an initial system minimization for 2000 steps and then simulated the system for 0.05 ns to allow the RNA to settle into the grid. After this initial equilibration, the process of unwinding was simulated by pulling on the strands through this grid, which acts as a tunnel, using the *distance colvars*, with a harmonic force. A soft restraining force was applied on the backbone of the nucleic acids. To avoid distortion of the double-helical shape of the RNA during pulling, the hydrogen bonds formed by the bases were enforced by virtual springs through the NAMD extraBonds feature, with a force of 20 kcal/mol/Å^2^.

### 5.2 FFEA simulations

#### 5.2.1 Meshing and construction of the RTC dimer for FFEA

Two representative meshes were constructed which correspond to the experimental data available; the cryo-EM density map from EMD-31138, a dimer form of the biologically active RTC, and the atomistic structure 7EGQ.pdb, which was used in the above NAMD atomistic molecular dynamics simulations. In the latter, the monomer RTC complex was formed from two separate meshes, one from the cryo-EM density map EMD-23008, corresponding to 7KRO.pdb, and the other from 7EGQ.pdb for the nsp14-10 complex atomistic information. These structures were used to generate 4 four interacting meshes; two in each monomer (Fig. 2). Meshes were constructed by first removing the dust and smoothing the mask using ChimeraX (Pettersen et al., 2021). The non-manifold elements within the surface were corrected with “MeshLab” (Cignoni et al., 2008), and course-grained with “MeshMixer” (Sommer et al., 2017). The 3D meshes were then generated by “Gmsh” (Geuzaine and Remacle, 2009) and made compatible for FFEA simulations with NetGen (Schöberl, 1997).

Following dust-removal and smoothing in ChimeraX, density for the single-stranded RNA was clearly visible in EMD-23008. However, this extremely thin structure is likely to cause instabilities in FFEA due to the small volumetric elements required to represent it. Therefore, the surface elements corresponding to RNA density were deleted, and the resulting surface repaired by filling with additional triangular elements in MeshMixer. The physical presence of the single-stranded RNA was then represented in the FFEA simulations using a harmonic spring.

#### 5.2.2 Visualizing FFEA trajectories using VMD

In order to create informative visualizations that contain multiple simulation modalities VMD was extended with new plugins to enable reading key FFEA simulation data. The new plugins enable VMD to read FFEA simulation and modeling data including tetrahedral mesh nodes, edges, faces, and topology information, as well as information about surface and interior faces and their properties, springs, and timevarying mesh node coordinates. The plugin extensions permit VMD to create visualizations showing the all-atom and FFEA simulations aligned and superimposed, with physical properties associated with the atoms, and FFEA nodes and faces available for use in assigning surface colors for graphical representations of the simulations.

### 5.3 AI-enabled multi-resolution simulations

We developed a hierarchy of AI methods that would facilitate embedding the high-dimensional (FFEA and AAMD) simulations into a low-dimensional manifold capturing both local (e.g., protein chain/subunit level) and global (e.g., mRTC monomer or dimer level) fluctuations. This required us to carefully consider the balance of simulation workloads and AI tools. We discuss the AI methods first, and then address the workload balance aspects in Sec. 5.4.

#### 5.3.1 Capturing global conformational transitions with anharmonic conformational analysis enabled autoencoders (ANCA-AE)

We have previously developed a hybrid machine learning approach that combines linear dimensionality reduction methods (Ramanathan et al., 2011) with a non-linear autoencoder. By minimizing the higher order (linear) correlations in the atomistic fluctuations and then examining non-linear correlations, we can obtain a succinct description of how global conformational changes are embodied in a simulation. This method, titled ANCA-AE (Clyde et al., 2021), can efficiently handle large dimensions (e.g., in the case of the mRTC, the 6,650 C*^α^*-atoms lead to approximately 20,000 dimensions) and can efficiently run on CPUs. We have shown that ANCA-AE can identify conformational states that share structural and energetic similarities, while characterizing transitional points in the high dimensional landscape. As shown in Fig. 5, we observed (for both FFEA and AAMD, although we only show results from FFEA simulations) that ANCA-AE extracts biophysically meaningful latent coordinates. The ANCA-AE method reveals conformational transitions from the FFEA trajectories, capturing the fluctuations in the interfaces between the nsp10-nsp14 complex and the rest of the RTC subcomponents.

**Figure 5:**
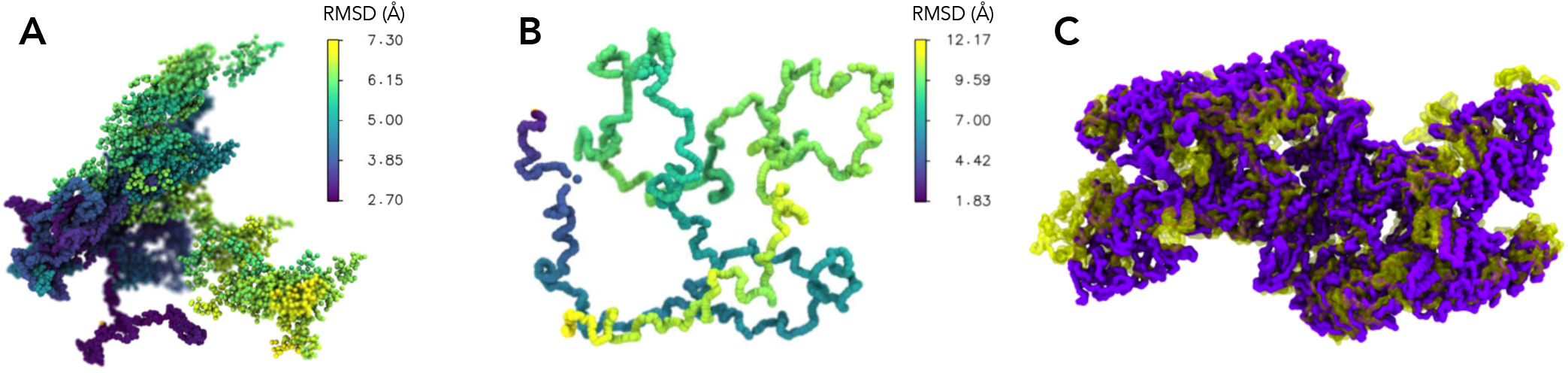
AI methods learn to summarize the time-dependent conformational changes in the FFEA and AAMD simulations. (A) The latent dimensional representation of the FFEA simulations summarized using the ANCA-AE method shows a clear separation between low-RMSD (deeper blue shades) and high-RMSD (yellow) states. (B) t-SNE visualization of the latent manifold learned using the graph neural operator (GNO) model on the entire RTC, painted by the system’s overall RMSD value. This model was trained on the entire 7egq system, comprised of 6650 total residues, with a window size of 10 frames. To offset the computational cost in the vast size of system, the model was trained on a small subset of the trajectory containing 2000 frames in order to investigate the method’s viability. The GNO has articulated the time-dependent conformational changes at the shorter timescales. (C) depicts the conformational changes between a low-RMSD and high-RMSD state learned from the GNO, capturing the relative motions of the nsp13 domains across the dimer interface.

#### 5.3.2 Describing localized conformational transitions with convolutional variational autoencoders (CVAE)

We also leveraged a deep learning algorithm, namely CVAE (Bhowmik et al., 2018) to describe localized conformational changes. The CVAE provides a compact representation of the high dimensional conformational landscape and can also capture intermediate/metastable states from long timescale MD simulations (Bhowmik et al., 2018). It can also be combined with outlier detection algorithms such as Density-Based Spatial Clustering of Applications with Noise (DBSCAN) (Ester et al., 1996) and the Local outlier factor (LOF) (Breunig et al., 2000) method to identify time points corresponding to rare transition events in the simulation trajectories. We note that both ANCA-AE and CVAE can utilize outlier detection algorithms for this purpose. Since we have previously discussed the CVAE model, we only present scaling results for this model.

However, CVAE is quadratic in time and space complexity (Casalino et al., 2021) and can be prohibitive to train for the entire mRTC. This makes the CVAE ideal to examine protein sub-unit fluctuations, which can be significantly smaller and more tractable for training. To optimize its performance, we implemented a distributed data parallel version of CVAE on the Summit supercomputer as well as a version within a single Cerebras CS-2 deep learning accelerator. By keeping all compute and memory resources on-silicon, the CS-2 provides, in a single system, orders of magnitude more compute power, on-chip memory, memory bandwidth, and communication bandwidth than traditional small-scale processors. This makes it particularly well-suited for accelerating ML models like the CVAE.

#### 5.3.3 Graph neural operator (GNO) networks for characterizing time-dependent conformational changes from AAMD simulations

Additionally, we also implemented the graph neural operator (GNO) as a fast surrogate model to simulate and predict the molecule movements from the AAMD simulation. The applications of deep learning for MD trajectory analysis, prediction and generation have recently exploded (Hoseini et al., 2021). The GNO model (Li et al., 2020) can solve partial differential equations (PDEs) by generalizing finite-dimensional neural networks to the infinite-dimensional operator-learning setting. In this work, we apply the GNO for the first time to capture the time dependent conformational changes governed by the system of equations in a traditional molecular dynamics simulation. As a first step, we show that we can reasonably predict protein backbone conformations up to 5 ps (250 MD simulation timesteps of 2 fs) into the future based on a 50 ps trajectory. This allows us to extract the time-dependent conformational changes of the molecule and analyze the connection between AAMD and FFEA simulation. This approach is capable of achieving an even longer prediction horizon of 500 ps on smaller test systems, and will be described in a separate manuscript. As shown in Fig. 5B, the ability to predict the time-dependent conformational changes in the entire RTC at short time-scales provides a means to accelerate sampling. The structure of the latent space following the RMSD transitions, provides insights into the large conformational changes captured in 5C.

### 5.4 Balsam workflow

Balsam (Salim et al., 2021) (Figure 6) provides a centralized service allowing users to register execution *sites* from any laptop, cluster, or supercomputer on which they wish to invoke computation. The sites run user agents that orchestrate workflows locally and periodically synchronize state with the central service. Users trigger remote computations by submitting jobs to the REST API, specifying the Balsam execution site and location(s) of data to be ingested.

**Figure 6:**
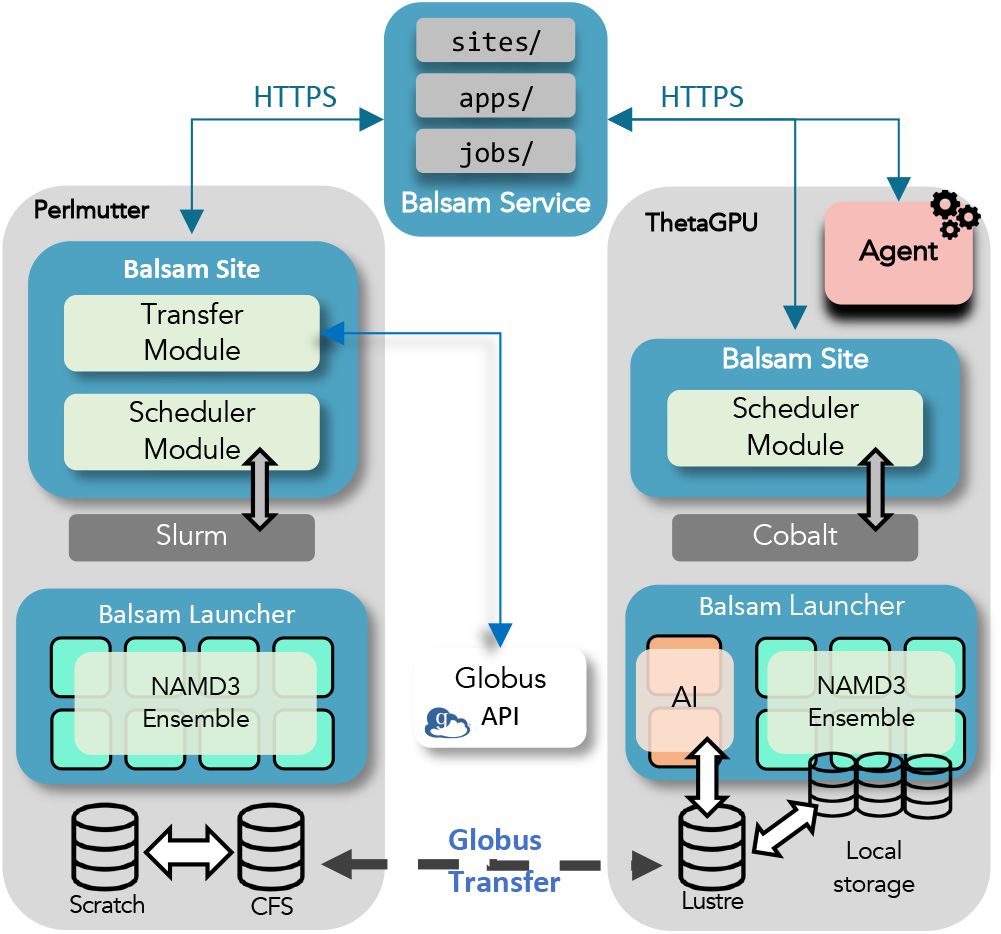
Cross-site Balsam Workflow architecture. Users execute scientific workloads by first creating Job resources at a specific execution site through the Balsam REST API. Pilot jobs at Balsam sites on the ALCF-ThetaGPU, and NERSC-Perlmutter supercomputers fetch the appropriate Jobs for execution. The MD Simulation and AI applications can span one or more sites based on the resource needs.

The ability to develop a unified workflow infrastructure that can span multiple computing sites by coupling traditional HPC simulation tasks with AI tasks is a key contribution of this paper. Further, the ability to steer a set of computational simulations based on AI and automatically switch simulation resolution (between allatom and continuum) is a further jump in capabilities.

A key development is the introduction of a central Balsam service that manages job distribution. Clients of this service use an API to inject and monitor jobs from anywhere. A second key development is the Balsam site, which consumes jobs from the Balsam service and interacts directly with the scheduling infrastructure of a cluster. By relying on a well-defined set of *platform interfaces* that adapt to machine-specific schedulers, node configurations, and pilot job launch paradigms, the Sites are easily portable across new and heterogeneous systems with minimal implementation effort. Together, the uniform Balsam API and portable Balsam Sites form a distributed, multi-resource execution landscape. The power of this approach was demonstrated using the Balsam API to inject simulation and AI jobs to the Balsam service, targeting simultaneous execution on multiple supercomputers. By strategically combining data transfer bundling and controlled job execution, Balsam achieves higher throughput than direct-scheduled local jobs and effectively balances job distribution across sites to simultaneously utilize multiple remote supercomputers.

The workflow driver agent comprised a lightweight Python process that used the Balsam SDK to track and dispatch NAMD 3 simulations on ThetaGPU^5^ and Perlmutter.^6^ A Python wrapper of a NAMD 3 replica’s execution enables the agent to seamlessly configure simulation parameters, dispatch runs, monitor data transfers, and send shutdown signals to simulations through a Python API. As simulation conformations arrive at the AI Site, the agent triggers model training and inference runs, whereby the outlier conformers are determined and used to initialize new simulations. In this scheme, a single Python agent exercises complete control over the geographically distributed simulation campaign. The workflow is robust to individual task- and batch job-level faults: a user monitoring the workflow with Balsam can reset failed jobs and provision additional resources by triggering new pilot jobs in the middle of a single agent’s running experiment. Experiments are launched with a simple YAML configuration format that flexibly supports arbitrarily many “simulation pools” on remote HPC systems: these pools comprise a particular simulation ensemble with data transfer settings (e.g. Globus transfer destination) which are passed through the Balsam Site’s Transfer module.

## 6 HOW PERFORMANCE WAS MEASURED

### 6.1 NAMD all-atom MD simulations

Production NAMD simulation performance is reported at runtime, directly yielding an achieved ns/day simulation rate. In order to report performance in terms of underlying machine FLOP rates, the marginal FLOPs per step are computed using hardware performance counters. Per-step FLOP performance metrics for NAMD 2.14 were collected on TACC Frontera^7^, using the Intel msr-tools utilities^8^, and the “TACC stats” system^9^. FLOP counts were measured for each NAMD simulation with runs of two different step counts. The results of the two different simulation lengths were subtracted to eliminate counting of startup-associated operations, to provide an accurate estimate of the FLOPs per step for a continuing simulation (Phillips et al., 2002).

### 6.2 AI methods

As outlined in Sec. 5.3, the ANCA-AE method is quite efficient and for the purposes of this paper, we did not need to optimize its performance. Further, we also note that the hybrid nature of combining linear- and non-linear dimensionality reduction opens up new avenues to assess scaling requirements from large datasets (which is beyond the scope of this paper). Similarly, the newly developed model, GNO was also not optimized for performance. We compared the performance of the GNO model with other baseline neural networks on a 28-node graph with 10,000 time frames. We report the total L2 error on a single GPU trained with 10 epochs.

We measure the throughput performance (samples/s) of the distributed training of CVAE model on the Summit supercomputer. The code is implemented in TensorFlow and parallelized with Horovod library in a data-parallel scheme. We then report the aggregated training throughput from 1 node up to 256 nodes (1536 GPUs). The scaling efficiency is determined by the ratio of ideal throughput (linear scaling) over the measured throughput. For results on the CS-2, we measured the throughput performance (samples/s) of training the CVAE model on a single Cerebras CS-2 system.

### 6.3 Balsam workflow

Balsam records each Job state transition (e.g. `RUNNING → RUN_DONE`) with timestamps and metadata that can be queried to produce workflow-level reports of task throughput and utilization (nodes occupied by running Balsam tasks, as a fraction of the currently provisioned resources). We then quantify the scientific throughput by measuring the cumulative MD simulation time, aggregated over all ensemble pools within an experiment). The Globus data transfer time is captured and recorded as part of the telemetry.

## 7 PERFORMANCE RESULTS

### 7.1 NAMD all-atom MD simulations

The performance of the NAMD production runs for the “RNA post-unwinding” are shown in Table 2, reporting CPU-based Frontera running NAMD 2 and two different DGX A100 320GB systems running GPU-resident NAMD 3. The first reported result was run on ThetaGPU, configured with one replica per GPU. The second reported result was run on NVIDIA Base Command,^10^ configured with one replica per node to simulate longer timescales. The Frontera and ThetaGPU measurements are averages. The median was used for NVIDIA Base Command jobs due to outliers; 3 of the 15 nodes achieved around 64ns/day, likely due to system I/O traffic.

NAMD 3 was benchmarked on the “RNA post-unwinding” system from Table 1 to characterize multi-GPU scaling performance. The equilibrated structure was run on a DGX A100 640GB node for 80000 timesteps using 8 CPU cores per GPU, and these simulations included the same parameters (e.g., 8Å cutoff) and file I/O as performed by the production runs, with trajectory output every 5000 steps. The results with estimated TFLOPS performance is reported in Table 3.

**Table 1:** Summary of AAMD simulations.

| Simulation | Non-equilibrium Sampling Methods | GFLOPs/step | Total sampling (μs) |
| --- | --- | --- | --- |
| Equilibration (pre-unwinding) | None | 24.9 | 2.107 |
| RNA unwinding | Extrabonds, Distance colvar, MDFF grid | 25.9 | 0.004 |
| RNA post-unwinding | Atom restraint, MDFF | 11.04 | 1.904 |
| RNA post-unwinding | None | 11.04 | 3.916 |

**Table 2:** NAMD “RNA post-unwinding” performance results.

| System | # of nodes per replica | ns/day |
| --- | --- | --- |
| TACC Frontera | 128 | 46.37 |
| NVIDIA DGX A100 320GB | 1/8 | 20.33 |
| NVIDIA DGX A100 320GB | 1 | 75.37 |

**Table 3:** NAMD 3 GPU-resident scaling on DGX A100 640GB.

| # of GPUs | ns/day | Scaling efficiency | Estimated TFLOPS |
| --- | --- | --- | --- |
| 1 | 20.7767 | 100% | 1.3274 |
| 2 | 36.1360 | 87% | 2.3087 |
| 4 | 56.3996 | 68% | 3.6033 |
| 8 | 75.2359 | 45% | 4.8067 |

The short-range non-bonded interaction calculations, which constitute most of the computational work per step, have long been optimized in NAMD by interpolating from a lookup table, effectively trading L1 cache reads for FLOPs. Doing this avoids expensive calculation of the error function `erf()` within the inner loop, as required by the particle-mesh Ewald (PME) method for calculating long-range electrostatics (Essmann et al., 1995). Use of a lookup table together with the CPU-based count of FLOPs per step leads to a very conservative estimate of TFLOPS for GPU-resident NAMD. GPU-based kernels actually produce a greater number of FLOPs per step than CPU-based kernels due to the need to accommodate a wider vector width. In NAMD 3, the non-bonded kernel computes interactions between tiles of 32 atoms, which allows for a high level of data reuse from on-chip memory while avoiding collisions and atomic operations. However, within a tile–tile calculation, a higher number of atom pairs will be outside of the cutoff radius, producing more GPU FLOPs than predicted by the CPU-based FLOP count.

As mentioned earlier, GPU scaling is primarily limited by the PME calculation. PME summation in reciprocal space requires computing 3D FFTs which cannot be decomposed efficiently across multiple GPUs, thus, the calculation is performed on a single GPU. Profiling the 8-GPU case shows that the GPU doing PME is overloaded at 80% utilization, while the rest of the GPUs are at 50% utilization. Strategies to mitigate this load imbalance include reducing the workload of the current PME implementation by distributing those parts of PME that are, in fact, scalable (in particular, the charge-spreading and force-interpolation procedures) and then employing task-based parallelism to reduce the amount of additional work assigned to the GPU doing the truly non-scalable PME work.

### 7.2 AI methods

As shown in Fig. 7, we achieved about 87% scaling efficiency in the distributed training of CVAE model with input size (512, 512) and (1024, 1024) on the Summit supercomputer. Equivalently, for an input size of 512 × 512, the CS-2 delivers out-of-the-box performance of 24,000 samples/s, or about the equivalent of 110-120 GPUs. For a larger input size of 1024 × 1024, CS-2 delivers 4,700 samples/s, also the equivalent of over 100 GPUs.

**Figure 7:**
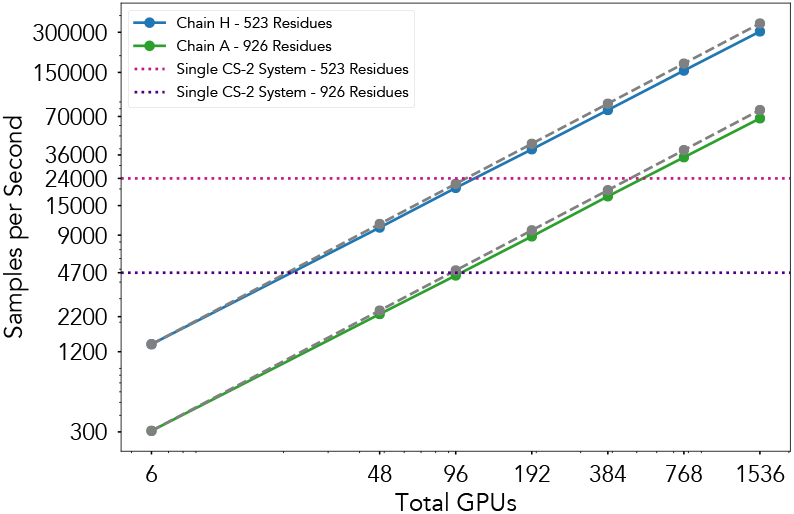
Scaling the CVAE model on Summit supercomputer (with increasing total GPUs) and the Cerebras CS-2 deep-learning accelerator. Chain H was cropped to a 512×512 contact map, and chain A was padded to a 1024×1024 contact map.

For these experiments, we measured performance as-is – without taking time to optimize the model or input functions for the Cerebras architecture. With additional time, we believe the performance from CS-2 would be much higher. Unlike with a typical distributed GPU setup, where hyperparameters need tuning as one scales from a single device to a large cluster of GPUs, the CS-2 is a single, powerful node. This means that we were able to run the CVAE model with the same experimental configuration as for a single GPU, without needing any hyperparameter changes.

Because a single CS-2 here delivers the performance of over 100 GPUs, it is a practical alternative for organizations interested in this workflow who do not have extremely large GPU clusters. By providing focused AI acceleration for the neural network portion of this application, the CS-2 also allows us to fully dedicate our GPU resources towards the more graphics-oriented simulation task.

### 7.3 Balsam workflow

We evaluate Balsam’s efficacy utilizing multiple HPC supercomputing sites to distribute the workloads and reduce the overall time to solution. In this experiment, the AAMD NAMD 3 simulations were run on both the Perlmutter and ThetaGPU systems. The AI training and inference models, in this case the ANCA-AE, were run on a dedicated ThetaGPU node. The trajectory information from the simulations is transferred via Globus from Perlmutter to ThetaGPU. This data together with the data from the local simulations at ThetaGPU are used to both re-train as well as to run inference. Based on the inference, promising simulation configurations are queued up with the Balsam service for launch and configurations that need to be stopped are terminated. We observe that balsam is able to sustain close to 100% utilization on Perlmutter (Fig. 8). The time taken by the AI models in this case can be completely overlapped with the simulations. We are also able to overlap the data transfer from Perlmutter to ThetaGPU. For wide-area Globus data transfers, we achieve a 44MB/sec mean effective bandwidth, with a wide distribution due to a standard deviation of 71 MB/sec. We attribute this to the shared network. However, we are able to overlap this data-movement and hide the latency. We are thus able to fully exploit distributed HPC resources spanning Perlmutter and ThetaGPU and able to accelerate the overall time to solution in comparison to executing at a single site.

**Figure 8:**
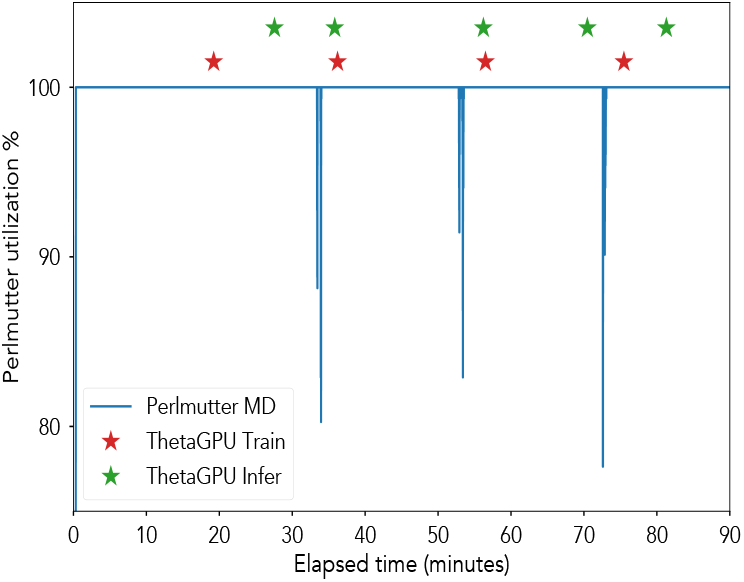
Cross-site Balsam Workflow spanning NERSC Perlmutter and ALCF ThetaGPU. We are able to achieve extreme utilization close to 100% on Perlmutter. Quick training and inference jobs, shown as stars on the timeline, are launched asynchronously on ThetaGPU upon receiving preprocessed training data from Perlmutter. The agent signals simulations to shutdown and restart from outlier conformations which leads to brief periods of inactivity on Perlmutter.

## 8 IMPLICATIONS

Our major scientific achievements include:

- Our studies reveal the inner workings of a large biomolecular machine, such as the SARS-CoV-2 RTC. By analyzing the fluctuations within the individual sub-components of this nano-machine, we were able to visualize how the viral-RNA is displaced to the respective active sites of the two key nsps.
- Although preliminary, our study reveals the concerted motions across the individual RTC monomers, which is coordinated by the interface proteins (nsp8, nsp9, nsp10). One of the nsp13 subunits on either side of the dimer also undergoes large fluctuations, displacing the bound RNA. This particular insight qualitatively agrees with the experimental data (Chen et al., 2020, Yan et al., 2020, 2021a); however provides a more detailed resolution of how such concerted motions are mediated across the entire RTC via protein-protein interfaces.
- The implicit coupling between AAMD and FFEA simulations by using our AI methods to learn interface potentials (that keep the protein-protein interfaces intact within the RTC simulations) reveal the dynamic nature of how large biomolecular assemblies.
- The capability to perform multi-site data analysis and simulations for integrative biology will be invaluable for making use of large experimental data that are difficult to transfer.
- We demonstrated the workflow across heterogeneous computational architectures, diverse HPC sites – two supercomputers, namely ThetaGPU (ALCF) and Perlmutter (NERSC). With the balkanization of heterogenous architecture and software platforms, the workflow represents a necessary capability that can be leveraged across domains where heterogeneous tasks and hardware platforms are the norm (for e.g., climate modeling, materials simulation, etc.). Further, our resource utilization is near 100% and optimally distributed for AI vs. HPC (simulation) jobs.
- The workflow infrastructure also illustrates some limitations in coordinating large file transfers across supercomputing sites that are required for both visualization and analysis.
- Our hierarchy of AI-methods – capturing local to global conformational changes, including the ability to predict time-dependent conformational changes can be extremely helpful in using AI to propagate MD / continuum simulations without having to resort to costly integration steps.

## ACKNOWLEDGMENTS

We thank the Argonne Leadership Computing Facility supported by the DOE under DE-AC02-06CH11357, the Oak Ridge Leadership Computing Facility at Oak Ridge National Laboratory supported by the DOE under Contract DE-AC05-00OR22725, and the National Energy Research Scientific Computing Center at Lawrence Berkeley National Laboratory supported by the DOE under Contract No. DE-AC02-05CH11231. We also thank the Texas Advanced Computing Center Frontera team, especially D. Stanzione and T. Cockerill, and for compute time made available through a Director’s Discretionary Allocation (NSF MCB-20024). NAMD and VMD are funded by NIH P41-GM104601. The NAMD team thanks Intel and M. Brown for contributing the AVX-512 tile list kernels. Anda Trifan acknowledges support from a DOE CSGF (DE-SC0019323). This research was supported by the Exascale Computing Project (17-SC-20-SC), a collaborative effort of the US DOE Office of Science and the National Nuclear Security Administration. Research was supported by the DOE through the National Virtual Biotechnology Laboratory, a consortium of DOE national laboratories focused on response to COVID-19, with funding from the Coronavirus CARES Act. This work used resources, services, and support from the COVID-19 HPC Consortium (https://covid19-hpc-consortium.org/), a privatepublic effort uniting government, industry, and academic leaders who are volunteering free compute time and resources in support of COVID-19 research.

## Footnotes

1 https://www.ks.uiuc.edu/Highlights/?section=2021&highlight=2021-09

2 https://www.mongodb.com/

3 https://api.nersc.gov/api/v1.2/

4 https://www.olcf.ornl.gov/summit/

5 https://www.alcf.anl.gov/support-center/theta/theta-thetagpu-overview

6 https://www.nersc.gov/systems/perlmutter/

7 https://www.tacc.utexas.edu/systems/frontera

8 https://github.com/intel/msr-tools

9 https://github.com/TACC/tacc_stats

10 https://www.nvidia.com/en-us/data-center/base-command-platform/

## Notes

### Competing Interest Statement

The authors have declared no competing interest.

## REFERENCES

M. L. Agostini, A. J. Pruijssers, J. D. Chappell, J. Gribble, X. Lu, E. L. Andres, G. R. Bluemling, M. A. Lockwood, T. P. Sheahan, A. C. Sims, et al. Small-molecule antiviral *β*-d-n 4-hydroxycytidine inhibits a proofreading-intact coronavirus with a high genetic barrier to resistance. Journal of virology, 93(24):e01348–19, 2019.

N. Alam and M. K. Higgins. A spike with which to beat covid-19? Nature Reviews Microbiology, 18(8):414–414, 2020. doi: 10.1038/s41579-020-0383-2. URL https://doi.org/10.1038/s41579-020-0383-2.

M. AlQuraishi. End-to-end differentiable learning of protein structure. Cell Systems, 8 (4):292 – 301.e3, 2019. ISSN 2405-4712. doi: https://doi.org/10.1016/j.cels.2019.03.006. URL http://www.sciencedirect.com/science/article/pii/S2405471219300766.

P. R. Arantes, A. Saha, and G. Palermo. Fighting covid-19 using molecular dynamics simulations. ACS Central Science, 6(10):1654–1656, 10 2020. doi: 10.1021/acscentsci.0c01236. URL https://doi.org/10.1021/acscentsci.0c01236.

V. Balasubramanian, A. Treikalis, O. Weidner, and S. Jha. Ensemble toolkit: Scalable and flexible execution of ensembles of tasks. In 2016 45th International Conference on Parallel Processing (ICPP), pages 458–463, Los Alamitos, CA, USA, aug 2016. IEEE Computer Society. doi: 10.1109/ICPP.2016.59. URL https://doi.ieeecomputersociety.org/10.1109/ICPP.2016.59.

M. Bárcena, C. O. Barnes, M. Beck, P. J. Bjorkman, B. Canard, G. F. Gao, Y. Gao, R. Hilgenfeld, G. Hummer, A. Patwardhan, G. Santoni, E. O. Saphire, C. Schaffitzel, S. L. Schendel, J. L. Smith, A. Thorn, D. Veesler, P. Zhang, and Q. Zhou. Structural biology in the fight against covid-19. Nature Structural & Molecular Biology, 28(1): 2–7, 2021. doi: 10.1038/s41594-020-00544-8. URL https://doi.org/10.1038/s41594-020-00544-8.

F. J. Barrantes. The contribution of biophysics and structural biology to current advances in covid-19. Annual Review of Biophysics, 50(1):493–523, 2021. doi: 10.1146/annurev-biophys-102620-080956. URL https://doi.org/10.1146/annurev-biophys-102620-080956. PMID: 33957057.

E. P. Barros, L. Casalino, Z. Gaieb, A. C. Dommer, Y. Wang, L. Fallon, L. Raguette, K. Belfon, C. Simmerling, and R. E. Amaro. The flexibility of ACE2 in the context of SARS-CoV-2 infection. bioRxiv, 2020. doi: 10.1101/2020.09.16.300459. URL https://www.biorxiv.org/content/early/2020/09/16/2020.09.16.300459.

D. Bhowmik, S. Gao, M. T. Young, and A. Ramanathan. Deep clustering of protein folding simulations. BMC Bioinformatics, 19(18):484, 2018. doi: 10.1186/s12859-018-2507-5. URL https://doi.org/10.1186/s12859-018-2507-5.

S. Bowerman, A. S. J. B. Rana, A. Rice, G. H. Pham, E. R. Strieter, and J. Wereszczynski. Determining atomistic saxs models of tri-ubiquitin chains from bayesian analysis of accelerated molecular dynamics simulations. Journal of Chemical Theory and Computation, 13(6):2418–2429, 06 2017. doi: 10.1021/acs.jctc.7b00059. URL https://doi.org/10.1021/acs.jctc.7b00059.

A. Brace, H. Lee, H. Ma, A. Trifan, M. Turilli, I. Yakushin, T. Munson, I. Foster, S. Jha, and A. Ramanathan. Achieving 100x faster simulations of complex biological phenomena by coupling ml to hpc ensembles. arXiv preprint arXiv:2104.04797, 2021.

L. A. Bratholm, A. S. Christensen, T. Hamelryck, and J. H. Jensen. Bayesian inference of protein structure from chemical shift data. PeerJ, 3:e861, 2015. ISSN 2167-8359 (Print); 2167-8359 (Electronic); 2167-8359 (Linking). doi: 10.7717/peerj.861.

M. M. Breunig, H.-P. Kriegel, R. T. Ng, and J. Sander. Lof: identifying density-based local outliers. In ACM sigmod record, volume 29, pages 93–104. ACM, 2000.

L. Casalino, A. C. Dommer, Z. Gaieb, E. P. Barros, T. Sztain, S.-H. Ahn, A. Trifan, A. Brace, A. T. Bogetti, A. Clyde, H. Ma, H. Lee, M. Turilli, S. Khalid, L. T. Chong, C. Simmerling, D. J. Hardy, J. D. Maia, J. C. Phillips, T. Kurth, A. C. Stern, L. Huang, J. D. McCalpin, M. Tatineni, T. Gibbs, J. E. Stone, S. Jha, A. Ramanathan, and R. E. Amaro. Ai-driven multiscale simulations illuminate mechanisms of sars-cov-2 spike dynamics. The International Journal of High Performance Computing Applications, 35(5):432–451, 2021. doi: 10.1177/10943420211006452. URL https://doi.org/10.1177/10943420211006452.

A. Cavalli, X. Salvatella, C. M. Dobson, and M. Vendruscolo. Protein structure determination from nmr chemical shifts. Proceedings of the National Academy of Sciences, 104(23):9615–9620, 2007. ISSN 0027-8424. doi: 10.1073/pnas.0610313104. URL https://www.pnas.org/content/104/23/9615.

R. Chard, Y. Babuji, Z. Li, T. Skluzacek, A. Woodard, B. Blaiszik, I. Foster, and K. Chard. funcx: A federated function serving fabric for science. ACM, Jun 2020. doi: 10.1145/3369583.3392683. URL http://dx.doi.org/10.1145/3369583.3392683.

J. Chen, B. Malone, E. Llewellyn, M. Grasso, P. M. Shelton, P. D. B. Olinares, K. Maruthi, E. T. Eng, H. Vatandaslar, B. T. Chait, T. M. Kapoor, S. A. Darst, and E. A. Campbell. Structural basis for helicase-polymerase coupling in the sars-cov-2 replication-transcription complex. Cell, 182(6):1560–1573.e13, 2020. ISSN 0092-8674. doi: 10.1016/j.cell.2020.07.033. URL https://www.sciencedirect.com/science/article/pii/S0092867420309417.

P. Cignoni, M. Callieri, M. Corsini, M. Dellepiane, F. Ganovelli, G. Ranzuglia, et al. Meshlab: an open-source mesh processing tool. In Eurographics Italian chapter conference, volume 2008, pages 129–136. Salerno, Italy, 2008.

A. Clyde, S. Galanie, D. W. Kneller, H. Ma, Y. Babuji, B. Blaiszik, A. Brace, T. Brettin, K. Chard, R. Chard, L. Coates, I. Foster, D. Hauner, V. Kertesz, N. Kumar, H. Lee, Z. Li, A. Merzky, J. G. Schmidt, L. Tan, M. Titov, A. Trifan, M. Turilli, H. Van Dam, S. C. Chennubhotla, S. Jha, A. Kovalevsky, A. Ramanathan, M. S. Head, and R. Stevens. High throughput virtual screening and validation of a sars-cov-2 main protease non-covalent inhibitor. bioRxiv, 2021. doi: 10.1101/2021.03.27.437323. URL https://www.biorxiv.org/content/early/2021/04/02/2021.03.27.437323.

F. A. Cruz and M. Martinasso. Firecrest: Restful api on cray xc systems, 2019.

D. De Sancho, J. A. Gavira, and R. Pérez-Jiménez. Coarse-grained molecular simulations of the binding of the sars-cov-2 spike protein rbd to the ace2 receptor. bioRxiv, 2020. doi: 10.1101/2020.05.07.083212. URL https://www.biorxiv.org/content/early/2020/07/24/2020.05.07.083212.

C. Dong and R. Skalak. Leukocyte deformability: Finite element modeling of large viscoelastic deformation. Journal of Theoretical Biology, 158(2):173–193, 1992. ISSN 0022-5193. doi: https://doi.org/10.1016/S0022-5193(05)80716-7. URL https://www.sciencedirect.com/science/article/pii/S0022519305807167.

A. E. P. Durumeric and G. A. Voth. Adversarial-residual-coarse-graining: Applying machine learning theory to systematic molecular coarse-graining. The Journal of Chemical Physics, 151(12):124110, 2019. doi: 10.1063/1.5097559. URL https://doi.org/10.1063/1.5097559.

T. M. Earnest, R. Watanabe, J. E. Stone, J. Mahamid, W. Baumeister, E. Villa, and Z. Luthey-Schulten. Challenges of integrating stochastic dynamics and cryoelectron tomograms in whole-cell simulations. The Journal of Physical Chemistry B, 121(15):3871–3881, 04 2017. doi: 10.1021/acs.jpcb.7b00672. URL https://doi.org/10.1021/acs.jpcb.7b00672.

P. Eastman, J. Swails, J. D. Chodera, R. T. McGibbon, Y. Zhao, K. A. Beauchamp, L.-P. Wang, A. C. Simmonett, M. P. Harrigan, C. D. Stern, et al. Openmm 7: Rapid development of high performance algorithms for molecular dynamics. PLoS computational biology, 13(7):e1005659, 2017. doi: 10.1371/journal.pcbi.1005659.

B. Enders, D. Bard, C. Snavely, L. Gerhardt, J. Lee, B. Totzke, K. Antypas, S. Byna, R. Cheema, S. Cholia, M. Day, A. Gaur, A. Greiner, T. Groves, M. Kiran, Q. Koziol, K. Rowland, C. Samuel, A. Selvarajan, A. Sim, D. Skinner, R. Thomas, and G. Torok. Cross-facility science with the superfacility project at lbnl. In 2020 IEEE/ACM 2nd Annual Workshop on Extreme-scale Experiment-in-the-Loop Computing (XLOOP), pages 1–7, Los Alamitos, CA, USA, nov 2020. IEEE Computer Society. doi: 10.1109/XLOOP51963.2020.00006. URL https://doi.ieeecomputersociety.org/10.1109/XLOOP51963.2020.00006.

B. D. Engel, M. Schaffer, L. Kuhn Cuellar, E. Villa, J. M. Plitzko, and W. Baumeister. Native architecture of the *Chlamydomonas* chloroplast revealed by in situ cryo-electron tomography. eLife, 4:e04889, jan 2015. ISSN 2050-084X. doi: 10.7554/eLife.04889. URL https://doi.org/10.7554/eLife.04889.

U. Essmann, L. Perera, M. L. Berkowitz, T. Darden, H. Lee, and L. G. Pedersen. A smooth particle mesh Ewald method. J. Chem. Phys., 103:8577–8593, 1995.

M. Ester, H.-P. Kriegel, J. Sander, and X. Xu. A density-based algorithm for discovering clusters in large spatial databases with noise. In Proc. of 2nd International Conference on Knowledge Discovery and Data Mining (KDD-96), pages 226–231, 1996.

G. Fiorin, M. L. Klein, and J. Hénin. Using collective variables to drive molecular dynamics simulations. Molecular Physics, 111(22-23):3345–3362, 2013. doi: 10.1080/00268976.2013.813594.

Y. Gao, L. Yan, Y. Huang, F. Liu, Y. Zhao, L. Cao, T. Wang, Q. Sun, Z. Ming, L. Zhang, et al. Structure of the rna-dependent rna polymerase from covid-19 virus. Science, 368(6492):779–782, 2020. doi: 10.1126/science.abb7498.

P. G. Garay, E. E. Barrera, F. Klein, M. R. Machado, M. Soñora, and S. Pantano. The sirah-cov-2 initiative: A coarse-grained simulations’ dataset of the sars-cov-2 proteome. Frontiers in Medical Technology, 3:4, 2021. ISSN 2673-3129. doi: 10.3389/fmedt.2021.644039. URL https://www.frontiersin.org/article/10.3389/fmedt.2021.644039.

C. Geuzaine and J.-F. Remacle. Gmsh: A 3-d finite element mesh generator with built-in pre-and post-processing facilities. International journal for numerical methods in engineering, 79(11):1309–1331, 2009.

A. Grishaev and M. Llinás. Protein structure elucidation from minimal nmr data: The clouds approach. In Nuclear Magnetic Resonance of Biological Macromolecules, volume 394 of Methods in Enzymology, pages 261 – 295. Academic Press, 2005. doi: 10.1016/S0076-6879(05)94010-X. URL http://www.sciencedirect.com/science/article/pii/S007668790594010X.

B. S. Hanson, S. Iida, D. J. Read, O. G. Harlen, G. Kurisu, H. Nakamura, and S. A. Harris. Continuum mechanical parameterisation of cytoplasmic dynein from atomistic simulation. Methods, 185:39–48, 2021.

P. Hoseini, L. Zhao, and A. Shehu. Generative deep learning for macromolecular structure and dynamics. Current Opinion in Structural Biology, 67:170–177, 2021.

W. Humphrey, A. Dalke, and K. Schulten. VMD – Visual Molecular Dynamics. J. Mol. Graphics, 14(1):33–38, 1996a. doi: 10.1016/0263-7855(96)00018-5.

W. Humphrey, A. Dalke, and K. Schulten. VMD: Visual molecular dynamics. Journal of Molecular Graphics, 1996b. ISSN 02637855. doi: 10.1016/0263-7855(96)00018-5.

B. E. Husic, N. E. Charron, D. Lemm, J. Wang, A. Pérez, A. Krämer, Y. Chen, S. Olsson, G. de Fabritiis, F. Noé, and C. Clementi. Coarse graining molecular dynamics with graph neural networks, 2020.

A. Jain, S. P. Ong, W. Chen, B. Medasani, X. Qu, M. Kocher, M. Brafman, G. Petretto, G.-M. Rignanese, G. Hautier, D. Gunter, and K. A. Persson. Fireworks: a dynamic workflow system designed for high-throughput applications. Concurrency and Computation: Practice and Experience, 27(17):5037–5059, 2015. ISSN 1532-0634. doi: 10.1002/cpe.3505. URL http://dx.doi.org/10.1002/cpe.3505.CPE-14-0307.R2.

J. Jumper, R. Evans, A. Pritzel, T. Green, M. Figurnov, O. Ronneberger, K. Tunyasuvunakool, R. Bates, A. Žídek, A. Potapenko, et al. Highly accurate protein structure prediction with alphafold. Nature, 596(7873):583–589, 2021. doi: 10.1038/s41586-021-03819-2.

L. Kalé, B. Acun, S. Bak, A. Becker, M. Bhandarkar, N. Bhat, A. Bhatele, E. Bohm, C. Bordage, R. Brunner, R. Buch, S. Chakravorty, K. Chandrasekar, J. Choi, M. Denardo, J. DeSouza, M. Diener, H. Dokania, I. Dooley, W. Fenton, J. Galvez, F. Gioachin, A. Gupta, G. Gupta, M. Gupta, A. Gursoy, V. Harsh, F. Hu, C. Huang, N. Jagathesan, N. Jain, P. Jetley, P. Jindal, R. Kanakagiri, G. Koenig, S. Krishnan, S. Kumar, D. Kunzman, M. Lang, A. Langer, O. Lawlor, C. Wai Lee, J. Lifflander, K. Mahesh, C. Mendes, H. Menon, C. Mei, E. Meneses, E. Mikida, P. Miller, R. Mokos, V. Narayanan, X. Ni, K. Nomura, S. Paranjpye, P. Ramachandran, B. Ramkumar, E. Ramos, M. Robson, N. Saboo, V. Saletore, O. Sarood, K. Senthil, N. Shah, W. Shu, A. B. Sinha, Y. Sun, Z. Sura, E. Totoni, K. Varadarajan, R. Venkataraman, J. Wang, L. Wesolowski, S. White, T. Wilmarth, J. Wright, J. Yelon, and G. Zheng. The Charm++ Parallel Programming System, Aug 2019. URL https://charm.cs.illinois.edu.

L. V. Kalé and G. Zheng. Chapter 1: The Charm++ Programming Model. In L. V. Kale and A. Bhatele, editors, Parallel Science and Engineering Applications: The Charm++ Approach, chapter 1, pages 1–16. CRC Press, Inc., Boca Raton, FL, USA, 1st edition, 2013. ISBN 1466504129, 9781466504127. doi: 10.1201/b16251.

R. Kennaway and E. Coen. Volumetric finite-element modelling of biological growth. Open Biology, 9(5):190057, 2019. doi: 10.1098/rsob.190057. URL https://royalsocietypublishing.org/doi/abs/10.1098/rsob.190057.

H. Kim and H. S. Jung. Cryo-em as a powerful tool for drug discovery: recent structural based studies of sars-cov-2. Applied Microscopy, 51(1):13, 2021. doi: 10.1186/s42649-021-00062-x. URL https://doi.org/10.1186/s42649-021-00062-x.

W. Kühlbrandt. The resolution revolution. Science, 343(6178):1443–1444, 2014. doi: 10.1126/science.1251652.

H. Lee, M. Turilli, S. Jha, D. Bhowmik, H. Ma, and A. Ramanathan. Deepdrivemd: Deep-learning driven adaptive molecular simulations for protein folding. In 2019 IEEE/ACM Third Workshop on Deep Learning on Supercomputers (DLS), pages 12–19, 2019.

Z. Li, N. Kovachki, K. Azizzadenesheli, B. Liu, K. Bhattacharya, A. Stuart, and A. Anandkumar. Neural operator: Graph kernel network for partial differential equations. arXiv preprint arXiv:2003.03485, 2020.

D. Lyumkis. Challenges and opportunities in cryo-em single-particle analysis. Journal of Biological Chemistry, 294(13):5181–5197, 2019. doi: 10.1074/jbc.REV118.005602.

J. Mahamid, S. Pfeffer, M. Schaffer, E. Villa, R. Danev, L. Kuhn Cuellar, F. Förster, A. A. Hyman, J. M. Plitzko, and W. Baumeister. Visualizing the molecular sociology at the hela cell nuclear periphery. Science, 351(6276):969–972, 2016. ISSN 0036-8075. doi: 10.1126/science.aad8857. URL https://science.sciencemag.org/content/351/6276/969.

A. Merk, A. Bartesaghi, S. Banerjee, V. Falconieri, P. Rao, M. I. Davis, R. Pragani, M. B. Boxer, L. A. Earl, J. L. Milne, and S. Subramaniam. Breaking cryo-em resolution barriers to facilitate drug discovery. Cell, 165(7):1698 – 1707, 2016. ISSN 0092-8674. doi: 10.1016/j.cell.2016.05.040. URL http://www.sciencedirect.com/science/article/pii/S0092867416305918.

A. Merzky, M. Turilli, M. Maldonado, M. Santcroos, and S. Jha. Using pilot systems to execute many task workloads on supercomputers, 2018.

B. Minkyung, D. Frank, A. Ivan, D. Justas, O. Sergey, L. G. Rie, W. Jue, C. Qian, K. L. N., S. R. Dustin, M. Claudia, P. Hahnbeom, A. Carson, G. C. R., D. Andy, P. J. H., R. A. V., van Dijk Alberdina A., E. A. C., O. D. J., S. Theo, B. Christoph, P.-K. Tea, R. M. K., D. Udit, Y. C. K., B. J. E., G. K. Christopher, G. N. V., A. P. D., R. R. J., and B. David. Accurate prediction of protein structures and interactions using a three-track neural network. Science, 373(6557):871–876, 2021/10/05 2021. doi: 10.1126/science.abj8754. URL https://doi.org/10.1126/science.abj8754.

E. N. Muratov, R. Amaro, C. H. Andrade, N. Brown, S. Ekins, D. Fourches, O. Isayev, D. Kozakov, J. L. Medina-Franco, K. M. Merz, T. I. Oprea, V. Poroikov, G. Schneider, M. H. Todd, A. Varnek, D. A. Winkler, A. V. Zakharov, A. Cherkasov, and A. Tropsha. A critical overview of computational approaches employed for covid-19 drug discovery. Chem. Soc.Rev., 50:9121–9151, 2021. doi: 10.1039/D0CS01065K. URL http://dx.doi.org/10.1039/D0CS01065K.

F. Noé. Machine Learning for Molecular Dynamics on Long Timescales, pages 331–372. Springer International Publishing, Cham, 2020. ISBN 978-3-030-40245-7. doi: 10.1007/978-3-030-40245-7_16. URL https://doi.org/10.1007/978-3-030-40245-7_16.

R. C. Oliver, D. J. Read, O. G. Harlen, and S. A. Harris. A stochastic finite element model for the dynamics of globular macromolecules. Journal of Computational Physics, 239:147–165, 2013. doi: 10.1016/j.jcp.2012.12.027.

A. K. Padhi, S. L. Rath, and T. Tripathi. Accelerating covid-19 research using molecular dynamics simulation. The Journal of Physical Chemistry B, 125(32):9078–9091, 08 2021. doi: 10.1021/acs.jpcb.1c04556. URL https://doi.org/10.1021/acs.jpcb.1c04556.

A. J. Pak and G. A. Voth. Advances in coarse-grained modeling of macromolecular complexes. Current Opinion in Structural Biology, 52:119–126, 2018. ISSN 0959-440X. doi: https://doi.org/10.1016/j.sbi.2018.11.005. URL https://www.sciencedirect.com/science/article/pii/S0959440X18300939. Cryo electron microscopy: the impact of the cryo-EM revolution in biology * Biophysical and computational methods - Part A.

O. Panagiotopoulou, J. Iriarte-Diaz, S. Wilshin, P. C. Dechow, A. B. Taylor, H. Mehari Abraha, S. F. Aljunid, and C. F. Ross. In vivo bone strain and finite element modeling of a rhesus macaque mandible during mastication. Zoology, 124:13–29, 2017. ISSN 0944-2006. doi: https://doi.org/10.1016/j.zool.2017.08.010. URL https://www.sciencedirect.com/science/article/pii/S0944200616301805. Determinants of Mammalian Feeding System Design.

J. K. Perry, T. C. Appleby, J. P. Bilello, J. Y. Feng, U. Schmitz, and E. A. Campbell. An atomistic model of the coronavirus replication-transcription complex as a hexamer assembled around nsp15. bioRxiv, 2021. doi: 10.1101/2021.06.08.447516. URL https://www.biorxiv.org/content/early/2021/06/08/2021.06.08.447516.

E. F. Pettersen, T. D. Goddard, C. C. Huang, E. C. Meng, G. S. Couch, T. I. Croll, J. H. Morris, and T. E. Ferrin. Ucsf chimerax: Structure visualization for researchers, educators, and developers. Protein Science, 30(1):70–82, 2021.

J. Phillips, G. Zheng, S. Kumar, and L. Kale. NAMD: Biomolecular simulation on thousands of processors. In Proceedings of the IEEE/ACM SC2002 Conference, Technical Paper 277, pages 1–18. IEEE Press, Baltimore, Maryland, 2002. doi: 10.1109/SC.2002.10019.

J. C. Phillips, R. Braun, W. Wang, J. Gumbart, E. Tajkhorshid, E. Villa, C. Chipot, R. D. Skeel, L. Kale, and K. Schulten. Scalable molecular dynamics with NAMD. J. Comput. Chem., 26:1781–1802, 2005.

J. C. Phillips, J. E. Stone, and K. Schulten. Adapting a message-driven parallel application to GPU-accelerated clusters. In SC ’08: Proceedings of the 2008 ACM/IEEE Conference on Supercomputing, pages 1–9, Piscataway, NJ, USA, 2008. IEEE Press. ISBN 978-1-4244-2835-9. (9 pages).

J. C. Phillips, D. J. Hardy, J. D. C. Maia, J. E. Stone, J. V. Ribeiro, R. C. Bernardi, R. Buch, G. Fiorin, J. Hénin, W. Jiang, R. McGreevy, M. C. R. Melo, B. Radak, R. D. Skeel, A. Singharoy, Y. Wang, B. Roux, A. Aksimentiev, Z. Luthey-Schulten, L. V. Kalé, K. Schulten, C. Chipot, and E. Tajkhorshid. Scalable molecular dynamics on CPU and GPU architectures with NAMD. J. Chem. Phys., 153:044130, 2020. doi: 10.1063/5.0014475.

A. Ramanathan, A. J. Savol, C. J. Langmead, P. K. Agarwal, and C. S. Chennubhotla. Discovering conformational sub-states relevant to protein function. PLOS ONE, 6 (1):1–16, 01 2011. doi: 10.1371/journal.pone.0015827. URL https://doi.org/10.1371/journal.pone.0015827.

R. A. Richardson, K. Papachristos, D. J. Read, O. G. Harlen, M. Harrison, E. Paci, S. P. Muench, and S. A. Harris. Understanding the apparent stator-rotor connections in the rotary atp ase family using coarse-grained computer modeling. Proteins: Structure, Function, and Bioinformatics, 82(12):3298–3311, 2014.

R. A. Richardson, B. S. Hanson, D. J. Read, O. G. Harlen, and S. A. Harris. Exploring the dynamics of flagellar dynein within the axoneme with fluctuating finite element analysis. Quarterly Reviews of Biophysics, 53, 2020.

M. Romano, A. Ruggiero, F. Squeglia, G. Maga, and R. Berisio. A structural view of sars-cov-2 rna replication machinery: Rna synthesis, proofreading and final capping. Cells, 9(5), 2020. ISSN 2073-4409. doi: 10.3390/cells9051267. URL https://www.mdpi.com/2073-4409/9/5/1267.

M. Salim, T. Uram, J. T. Childers, V. Vishwanath, and M. E. Papka. Toward realtime analysis of experimental science workloads on geographically distributed supercomputers, 2021.

S. H. Scheres. A bayesian view on cryo-em structure determination. Journal of Molecular Biology, 415(2):406 – 418, 2012. ISSN 0022-2836. doi: https://doi.org/10.1016/j.jmb.2011.11.010. URL http://www.sciencedirect.com/science/article/pii/S0022283611012290.

J. Schöberl. Netgen an advancing front 2d/3d-mesh generator based on abstract rules. Computing and visualization in science, 1(1):41–52, 1997.

M. Sener, S. Levy, J. E. Stone, A. Christensen, B. Isralewitz, R. Patterson, K. Borkiewicz, J. Carpenter, C. N. Hunter, Z. Luthey-Schulten, and D. Cox. Multiscale modeling and cinematic visualization of photosynthetic energy conversion processes from electronic to cell scales. Parallel Computing, page 102698, 2021.

J. Shang, Y. Wan, C. Luo, G. Ye, Q. Geng, A. Auerbach, and F. Li. Cell entry mechanisms of sars-cov-2. Proceedings of the National Academy of Sciences, 117(21):11727–11734, 2020.

T. P. Sheahan, A. C. Sims, S. Zhou, R. L. Graham, A. J. Pruijssers, M. L. Agostini, S. R. Leist, A. Schäfer, K. H. Dinnon, L. J. Stevens, et al. An orally bioavailable broadspectrum antiviral inhibits sars-cov-2 in human airway epithelial cell cultures and multiple coronaviruses in mice. Science translational medicine, 12(541), 2020.

A. Solernou, B.S. Hanson, R. A. Richardson, R. Welch, D.J. Read, O. G. Harlen, and S. A. Harris. Fluctuating finite element analysis (ffea): A continuum mechanics software tool for mesoscale simulation of biomolecules. PLoS computational biology, 14(3): e1005897, 2018.

K. Sommer, R. L. Izzo, L. Shepard, A. R. Podgorsak, S. Rudin, A. H. Siddiqui, M. F. Wilson, E. Angel, Z. Said, M. Springer, et al. Design optimization for accurate flow simulations in 3d printed vascular phantoms derived from computed tomography angiography. In Medical Imaging 2017: Imaging Informatics for Healthcare, Research, and Applications, volume 10138, page 101380R. International Society for Optics and Photonics, 2017.

J. E. Stone, B. Isralewitz, and K. Schulten. Early experiences scaling VMD molecular visualization and analysis jobs on Blue Waters. In Extreme Scaling Workshop (XSW), 2013, pages 43–50, Aug. 2013a. doi: 10.1109/XSW.2013.10.

J. E. Stone, K. L. Vandivort, and K. Schulten. GPU-accelerated molecular visualization on petascale supercomputing platforms. In Proceedings of the 8th International Workshop on Ultrascale Visualization, UltraVis ’13, pages 6:1–6:8, New York, NY, USA, 2013b. ACM.

J. E. Stone, R. McGreevy, B. Isralewitz, and K. Schulten. GPU-accelerated analysis and visualization of large structures solved by molecular dynamics flexible fitting. Faraday Discussions, 169:265–283, 2014. doi: 10.1039/C4FD00005F.

J. E. Stone, M. Sener, K. L. Vandivort, A. Barragan, A. Singharoy, I. Teo, J. V. Ribeiro, B. Isralewitz, B. Liu, B. C. Goh, J. C. Phillips, C. MacGregor-Chatwin, M. P. Johnson, L. F. Kourkoutis, C. N. Hunter, and K. Schulten. Atomic detail visualization of photosynthetic membranes with GPU-accelerated ray tracing. Parallel Computing, 55:17–27, 2016. doi: 10.1016/j.parco.2015.10.015.

T. Sztain, S.-H. Ahn, A. T. Bogetti, L. Casalino, J. A. Goldsmith, E. Seitz, R. S. McCool, F. L. Kearns, F. Acosta-Reyes, S. Maji, et al. A glycan gate controls opening of the sars-cov-2 spike protein. Nature Chemistry, pages 1–6, 2021.

V. Tozzini. Multiscale modeling of proteins. Accounts of Chemical Research, 43(2): 220–230, 02 2010. doi: 10.1021/ar9001476. URL https://doi.org/10.1021/ar9001476.

L. G. Trabuco, E. Villa, E. Schreiner, C. B. Harrison, and K. Schulten. Molecular dynamics flexible fitting: a practical guide to combine cryo-electron microscopy and x-ray crystallography. Methods, 49(2):174–180, 2009.

K. Tunyasuvunakool, J. Adler, Z. Wu, T. Green, M. Zielinski, A. Žídek, A. Bridgland, A. Cowie, C. Meyer, A. Laydon, S. Velankar, G. J. Kleywegt, A. Bateman, R. Evans, A. Pritzel, M. Figurnov, O. Ronneberger, R. Bates, S. A. A. Kohl, A. Potapenko, A. J. Ballard, B. Romera-Paredes, S. Nikolov, R. Jain, E. Clancy, D. Reiman, S. Petersen, A. W. Senior, K. Kavukcuoglu, E. Birney, P. Kohli, J. Jumper, and D. Hassabis. Highly accurate protein structure prediction for the human proteome. Nature, 596(7873): 590–596, 2021. doi: 10.1038/s41586-021-03828-1. URL https://doi.org/10.1038/s41586-021-03828-1.

T. W. van der Heijden, D. J. Read, O. G. Harlen, P. van der Schoot, S. A. Harris, and C. Storm. Combined force-torque spectroscopy of proteins by means of multiscale molecular simulation. Biophysical Journal, 119(11):2240–2250, 2020.

K. Vanommeslaeghe, E. Hatcher, C. Acharya, S. Kundu, S. Zhong, J. Shim, E. Darian, O. Guvench, P. Lopes, I. Vorobyov, et al. Charmm general force field: A force field for drug-like molecules compatible with the charmm all-atom additive biological force fields. Journal of computational chemistry, 31(4):671–690, 2010.

J. W. Vant, S.-L. J. Lahey, K. Jana, M. Shekhar, D. Sarkar, B. H. Munk, U. Kleinekathofer, S. Mittal, C. Rowley, and A. Singharoy. Flexible fitting of small molecules into electron microscopy maps using molecular dynamics simulations with neural network potentials. Journal of chemical information and modeling, 60(5):2591–2604, 2020.

E. Villa and K. Lasker. Finding the right fit: chiseling structures out of cryoelectron microscopy maps. Current Opinion in Structural Biology, 25:118 – 125, 2014. ISSN 0959-440X. doi: https://doi.org/10.1016/j.sbi.2014.04.001. URL http://www.sciencedirect.com/science/article/pii/S0959440X14000414. Theory and simulation / Macromolecular machines.

J. Walpole, J. A. Papin, and S. M. Peirce. Multiscale computational models of complex biological systems. Annual Review of Biomedical Engineering, 15(1):137–154, 2013. doi: 10.1146/annurev-bioeng-071811-150104. URL https://doi.org/10.1146/annurev-bioeng-071811-150104. PMID: 23642247.

Q. Wang, J. Wu, H. Wang, Y. Gao, Q. Liu, A. Mu, W. Ji, L. Yan, Y. Zhu, C. Zhu, et al. Structural basis for rna replication by the sars-cov-2 polymerase. Cell, 182(2): 417–428, 2020.

D. B. Wells, V. Abramkina, and A. Aksimentiev. Exploring transmembrane transport through *α*-hemolysin with grid-steered molecular dynamics. The Journal of chemical physics, 127(12):09B619, 2007.

A. Wu, Y. Peng, B. Huang, X. Ding, X. Wang, P. Niu, J. Meng, Z. Zhu, Z. Zhang, J. Wang, et al. Genome composition and divergence of the novel coronavirus (2019-ncov) originating in china. Cell host & microbe, 27(3):325–328, 2020.

L. Yan, Y. Zhang, J. Ge, L. Zheng, Y. Gao, T. Wang, Z. Jia, H. Wang, Y. Huang, M. Li, Q. Wang, Z. Rao, and Z. Lou. Architecture of a sars-cov-2 mini replication and transcription complex. Nature Communications, 11(1):5874, 2020. doi: 10.1038/s41467-020-19770-1. URL https://doi.org/10.1038/s41467-020-19770-1.

L. Yan, J. Ge, L. Zheng, Y. Zhang, Y. Gao, T. Wang, Y. Huang, Y. Yang, S. Gao, M. Li, Z. Liu, H. Wang, Y. Li, Y. Chen, L. W. Guddat, Q. Wang, Z. Rao, and Z. Lou. Cryo-em structure of an extended sars-cov-2 replication and transcription complex reveals an intermediate state in cap synthesis. Cell, 184(1):184–193.e10, 2021a. ISSN 0092-8674. doi: https://doi.org/10.1016/j.cell.2020.11.016. URL https://www.sciencedirect.com/science/article/pii/S0092867420315336.

L. Yan, Y. Yang, M. Li, Y. Zhang, L. Zheng, J. Ge, Y. C. Huang, Z. Liu, T. Wang, S. Gao, et al. Coupling of n7-methyltransferase and 3-5 exoribonuclease with sars-cov-2 polymerase reveals mechanisms for capping and proofreading. Cell, 184(13): 3474–3485, 2021b.

A. Yu, A. J. Pak, P. He, V. Monje-Galvan, L. Casalino, Z. Gaieb, A. C. Dommer, R. E. Amaro, and G. A. Voth. A multiscale coarse-grained model of the sars-cov-2 virion. Biophysical Journal, 120(6):1097–1104, 2021. ISSN 0006-3495. doi: https://doi.org/10.1016/j.bpj.2020.10.048. URL https://www.sciencedirect.com/science/article/pii/S0006349520331684.

Q. Zhang, R. Xiang, S. Huo, Y. Zhou, S. Jiang, Q. Wang, and F. Yu. Molecular mechanism of interaction between sars-cov-2 and host cells and interventional therapy. Signal Transduction and Targeted Therapy, 6(1):1–19, 2021.

H.-X. Zhou. Theoretical frameworks for multiscale modeling and simulation. Current Opinion in Structural Biology, 25:67–76, 2014. doi: https://doi.org/10.1016/j.sbi.2014.01.004. URL https://www.sciencedirect.com/science/article/pii/S0959440X14000050.

M. I. Zimmerman, J. R. Porter, M. D. Ward, S. Singh, N. Vithani, A. Meller, U. L. Mallimadugula, C. E. Kuhn, J. H. Borowsky, R. P. Wiewiora, M. F. D. Hurley, A. M. Harbison, C. A. Fogarty, J. E. Coffland, E. Fadda, V. A. Voelz, J. D. Chodera, and G. R. Bowman. Sars-cov-2 simulations go exascale to capture spike opening and reveal cryptic pockets across the proteome. bioRxiv, 2020. doi: 10.1101/2020.06.27.175430. URL https://www.biorxiv.org/content/early/2020/10/07/2020.06.27.175430.

